# Protocol for a minigene splice assay using the pET01 vector

**DOI:** 10.1101/2025.02.03.636202

**Authors:** Hannah Andreae, Marialessandra Curcio, Sahar Esmaeelpour, Friederike Jahnke, Fritz Benseler, Nils Brose, Barbara Vona

**Affiliations:** Institute of Human Genetics, University Medical Center Göttingen, 37073 Göttingen, Germany; Institute for Auditory Neuroscience and InnerEarLab, University Medical Center Göttingen, 37073 Göttingen, Germany; Department of Molecular Neurobiology, Max Planck Institute for Multidisciplinary Sciences, 37075 Göttingen, Germany

## Abstract

Aberrant splicing is recognized as a key contributor of hereditary disorders, yet characterizing the molecular effects of splice variants is an important task that poses several challenges. Here, we present a protocol for an *in vitro* splice assay using a minigene approach, which is especially useful when patient samples are not available for RNA analysis or when target genes or isoforms are not expressed in accessible tissues for direct analysis. We describe steps for assay design, including the cloning of minigene plasmids and subsequent transfection, followed by RNA isolation and cDNA synthesis. We also provide details on quantitative fragment analysis using capillary electrophoresis and optional subcloning to facilitate optimal sequencing of multiple splicing products.

## Before you begin

An important effect of nucleotide sequence variations is their potential to significantly alter the splicing signal landscape utilized by the spliceosome. This is particularly relevant when analyses are solely conducted at the DNA level without incorporating RNA studies.^1,2^

In our investigation of genetic variants in otoferlin (*OTOF*), a protein expressed exclusively in the cochlea and brain, we found no practical methods to characterize splicing effects using commonly available patient tissues, such as blood or fibroblasts.^3^ We describe here an *in vitro* minigene splice assay that resolves this issue and provides an effective approach to elucidate spliceogenic effects in the absence of target tissues. Based on this method, we present data for four *OTOF* variants reported in patients, detailed in the expected results section and in supplementary material.

This protocol outlines the materials and methods required to functionally characterize genetic variants predicted to cause splicing aberrations using an *in vitro* (minigene) assay with the exon-trapping vector pET01 (MoBiTec GmbH, Göttingen, Germany). This vector contains an intrinsic splicing function, comprising a promoter followed by exon A, an intron with a multiple cloning site (into which the variant-containing region is cloned), exon B, and a sequence for the Poly-A tail (Figure 1). Upon transfection into eukaryotic cells, RNA expression and splicing of the mature RNA occurs (see datasheet “Exontrap” by MoBiTec GmbH^4^), allowing assessment of the variant impact on splicing that can be compared to the wild-type sequence.

**Figure 1:**
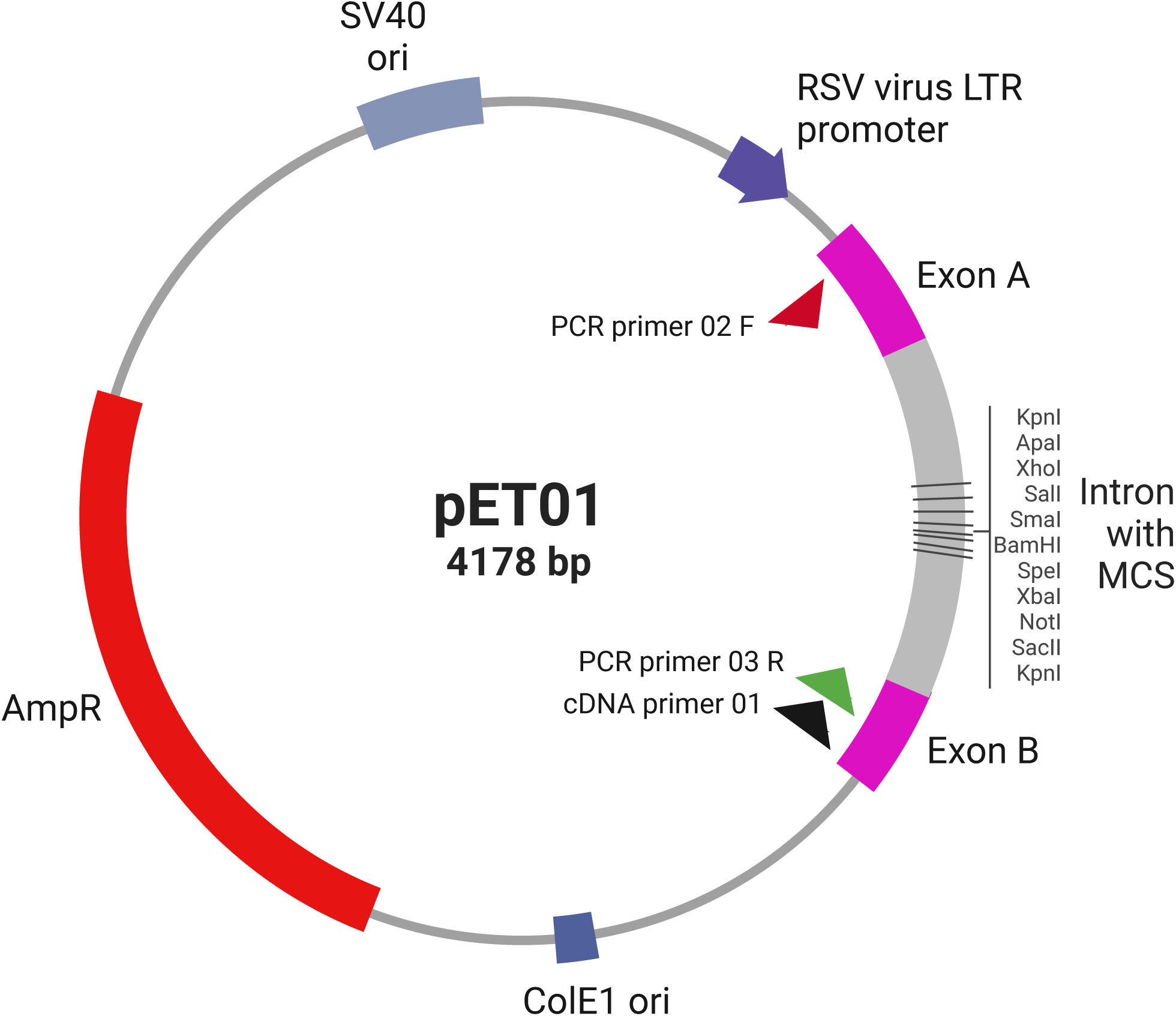
pET01 plasmid map illustrating the multiple cloning site located within the intron, flanked by exons A and B, along with the primers used in this protocol aligned to the vector (adapted from datasheet “Exontrap” by MoBiTec GmbH^4^).

In summary, we present an approach for a splice assay that facilitates the investigation of splice effects of genetic variants when patient RNA is unavailable or when gene expression of target genes or isoforms is absent in accessible tissues for direct analysis.

### Institutional permissions (if applicable)

Before beginning this protocol, users must ensure that informed consent and ethical approval have been obtained for the use of patient-derived DNA. Institutional permission must be secured, if applicable. The example shown here uses DNA from a healthy volunteer, thus negating the need for ethical approval.

### Design assay-specific primers

#### Timing: Variable

Specific primers are necessary for amplifying the region containing the variant of interest, thereby creating the insert for pET01. In cases where patient DNA with the variant of interest is unavailable, primers for site-directed mutagenesis are required.

1. Design primers to amplify a large target region for cloning into pET01. The following steps a-f exemplify a standard procedure for primer design.

a. Select one or more exons, ensuring at least 60 base pairs (bp) of flanking introns. We usually work with inserts ranging from 500-700 bp or more. The DNA sequence can be retrieved using Ensembl (www.ensembl.org/).
b. The selected sequence is used as a source sequence in Primer3web (https://primer3.ut.ee). Set the “Primer GC percentage” to a minimum of 47%, optimal at 50% and maximum of 53%. Select appropriate primers.
c. Use SNPCheck (https://genetools.org/SNPCheck/snpcheck.htm) to detect single nucleotide polymorphisms (SNPs) ensuring optimal primer binding. SNPs should be minimized or have a low minor allele frequency (e.g. <0.01).
d. Perform multiple alignment queries against the human genome using UCSC Genome Browser Human BLAT Search tool (https://genome.ucsc.edu/cgi-bin/hgBlat).
e. Identify two different restriction enzymes to use based on the multiple cloning sites (refer to the multiple cloning site of pET01 “Exontrap Handbook”^4^ or Figure 1) and incorporate the corresponding sites with three to four additional nucleotides at the 5’ end of the primers in the correct orientation.
f. Confirm that no restriction sites are present within the insert sequence using the ApE (A plasmid Editor) application (https://jorgensen.biology.utah.edu/wayned/ape/).

**Critical:** Select two appropriate different restriction sites to be added to the forward and reverse primer respectively (e.g. ApaI, XhoI, SalI, SmaI, BamHI, SpeI, XbaI, NotI, SacII) to ensure correct orientation of the insert in the pET01 vector.

2. Design primers for site-directed mutagenesis with the NEBaseChanger online tool (https://nebasechanger.neb.com/).

### Other preparations

#### Timing: Variable

1. Ensure that the desired cell line for transfection is seeded into a 6-well plate at a density of 2 × 10^5^ cells/ml (approximately 400,000 cells per well) the day prior to transfection, accounting for the time required to achieve confluency in culture prior to seeding.

2. Obtain the pET01 vector in advance.

a. Rehydrate the dried vector according to the manufacturer’s instructions.
b. Transform the vector into *E. coli* according to standard lab protocols.
c. Plate on an ampicillin-containing LB agar plate and incubate overnight at 37°C.
d. Pick a colony to inoculate an overnight culture with ampicillin, incubating overnight with shaking. **Critical:** Make sure to store back-up overnight culture as a glycerol stock in -80°C.
e. Perform a standard plasmid isolation and elute the pET01 vector with dH_2_O.

**Note:** For a comprehensive overview of pET01 we provide the full plasmid sequence with already marked primer and restriction sites. Download the document under S1_pET01_ sequence.

## Key resources table

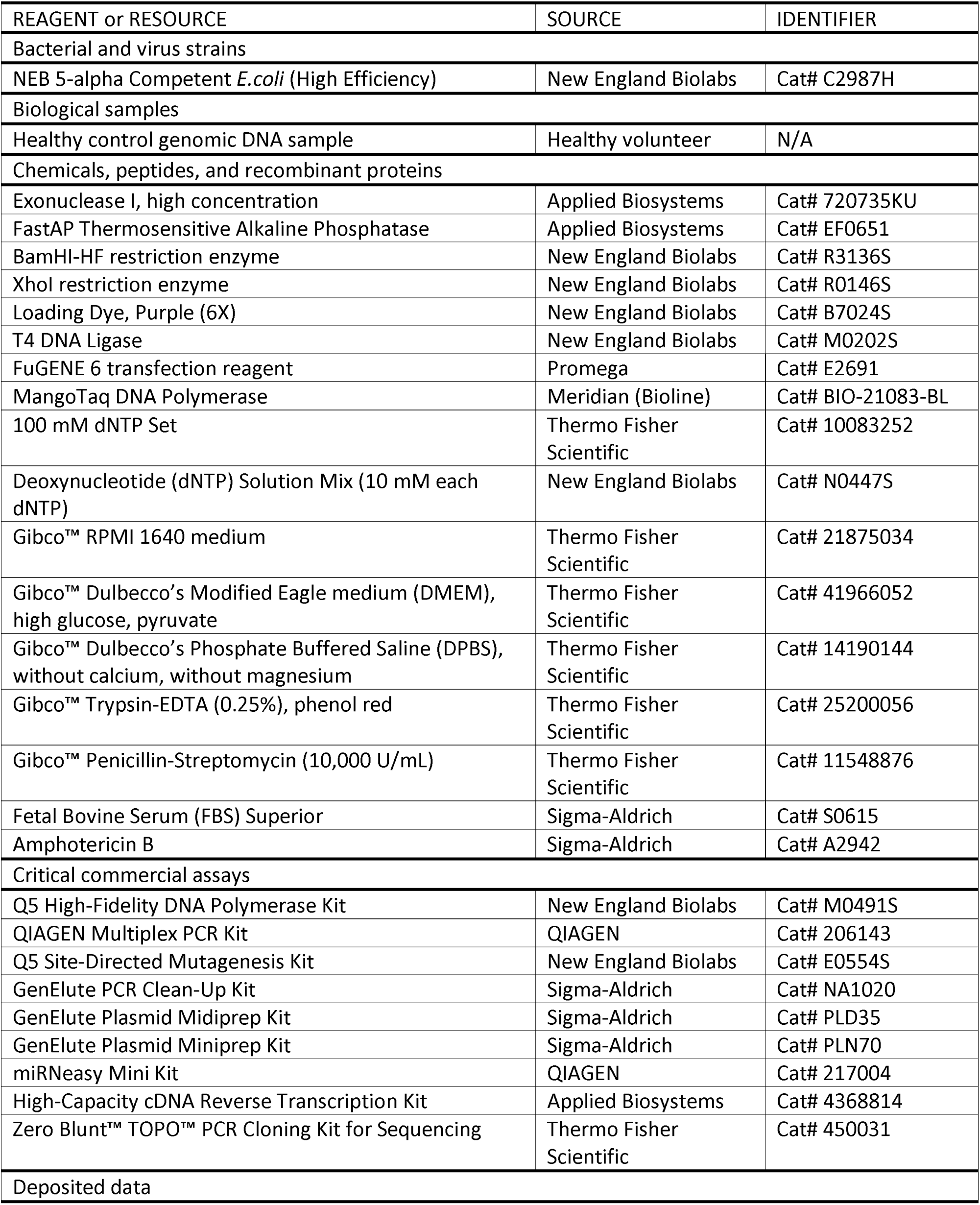
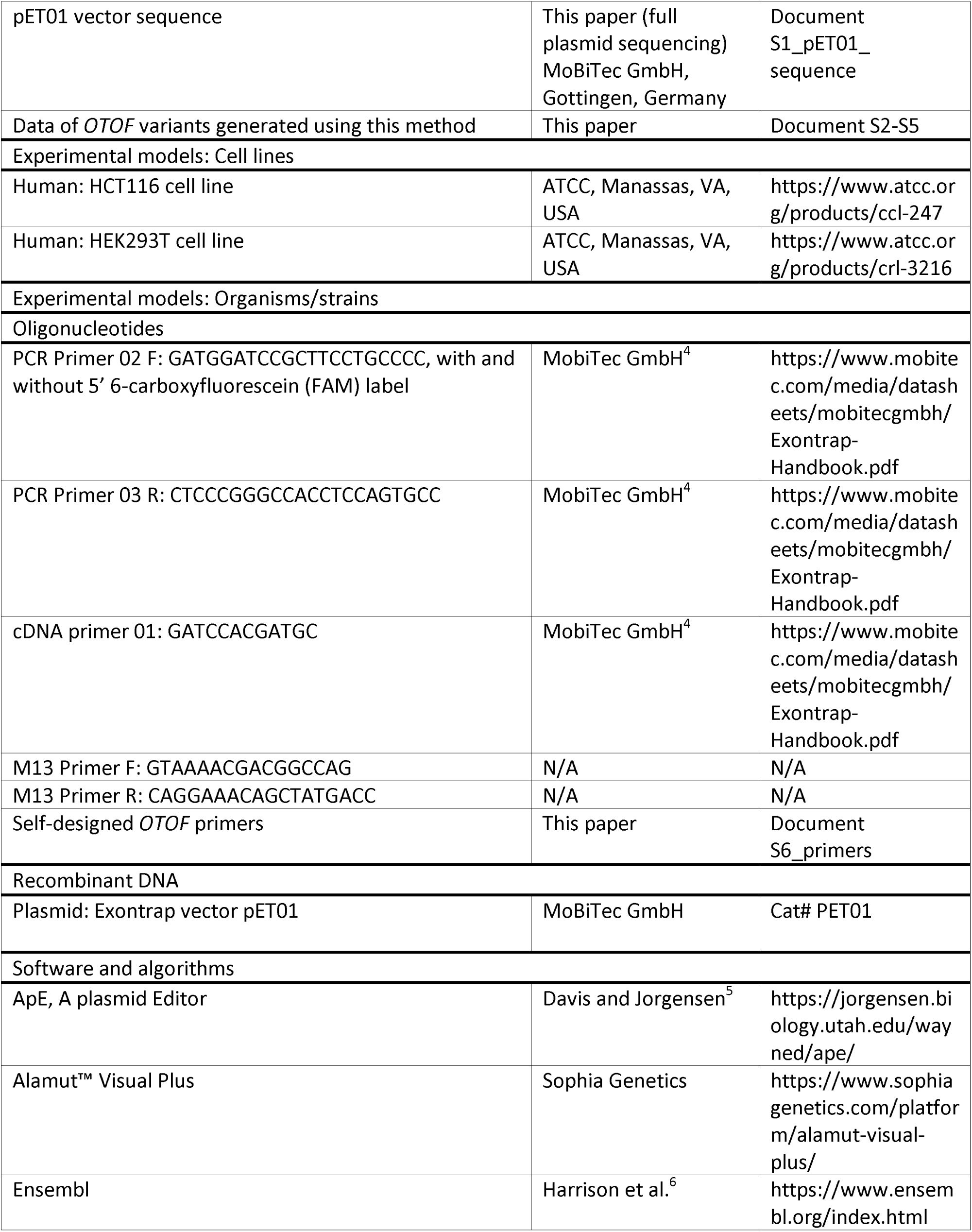
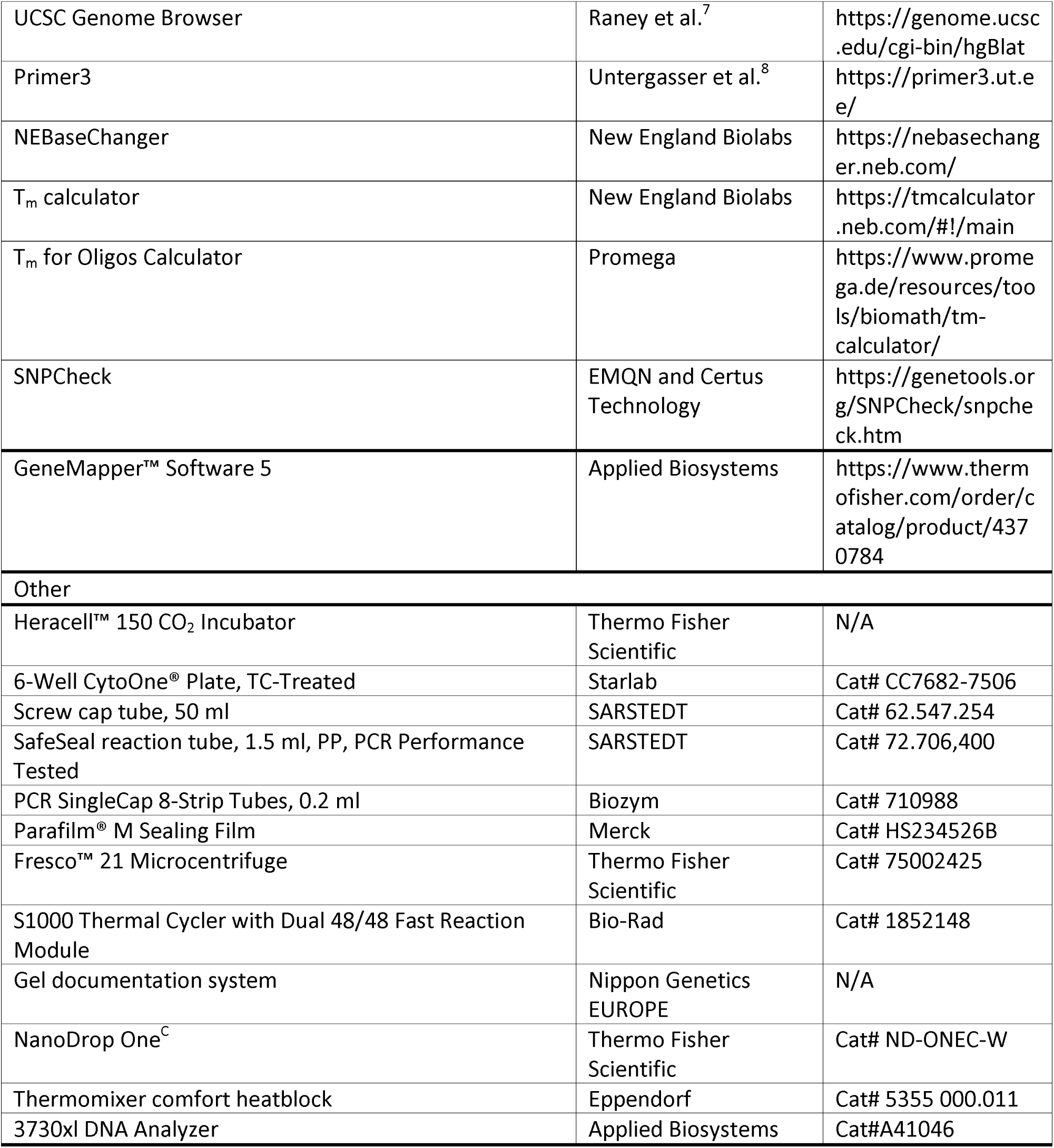

## Materials and equipment setup (optional)

### Supplemented RPMI/DMEM medium

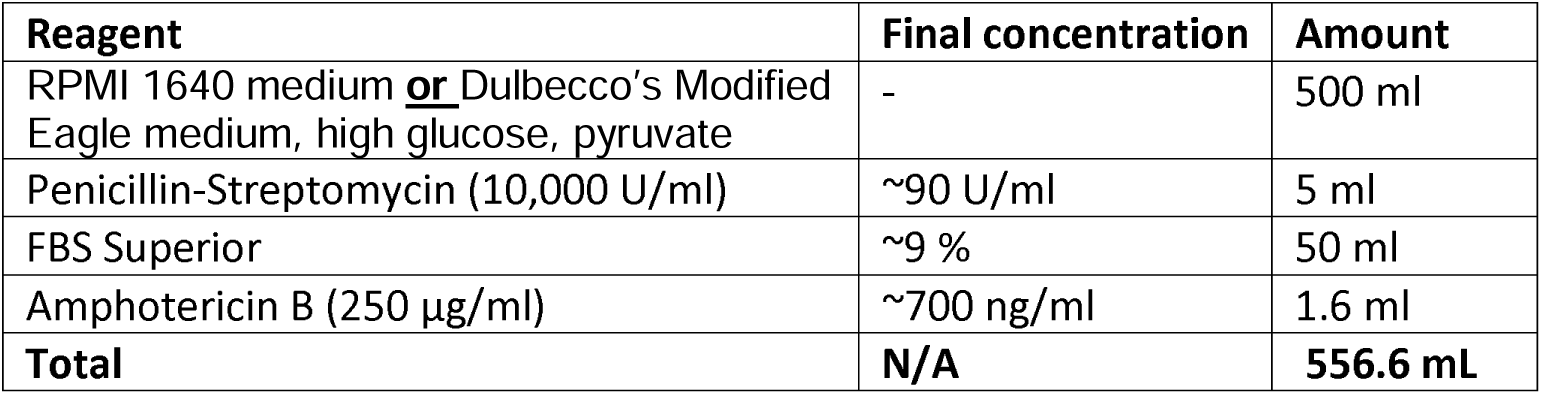

**Note:** Store at 4°C for up to 1 month and pre-warm to 37°C before usage.

### LB agar plates with selection marker (50 µg/ml)

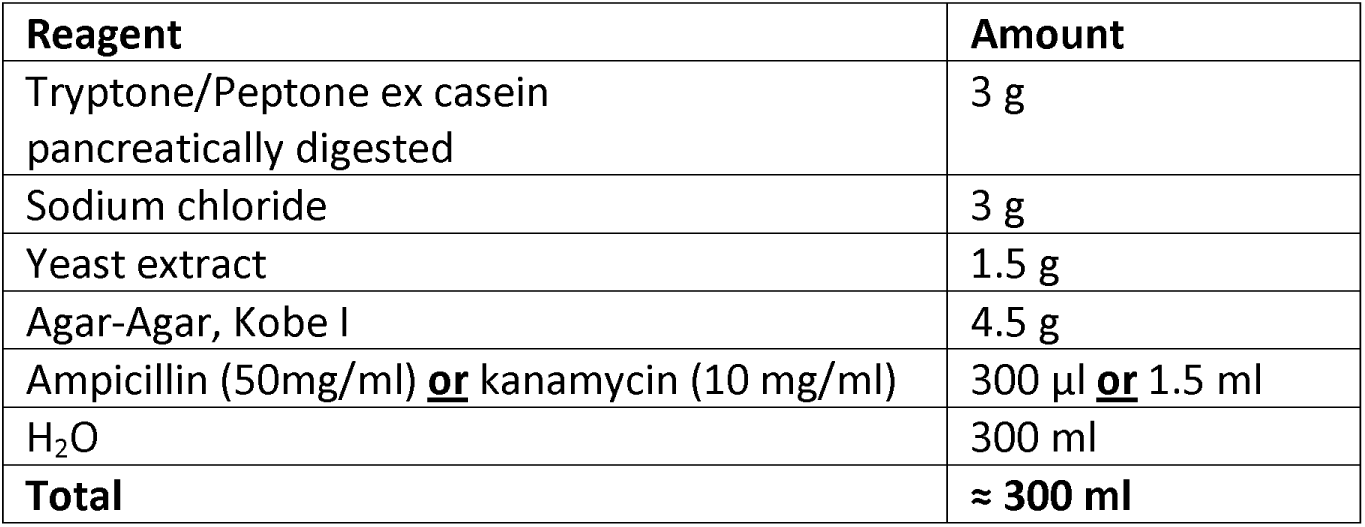

**Note:** Autoclave the LB agar mixture and let it cool down to ∼60°C before adding the antibiotics. Then pour into petri dishes and let them solidify. Store at 4°C.

## Step-by-step method details

### Amplify your region of interest

#### Timing: 2-3 h

1. Determine PCR conditions.

a. Calculate the annealing temperature of your designed primers according to the manufacturer’s instructions. We use the NEB T_m_ Calculator website (https://tmcalculator.neb.com/#!/main) recommended for the Q5 High-Fidelity DNA Polymerase Kit with the primer concentration set to 500 nM.
b. Calculate the extension time of the PCR. We use 25 s per kilobase of amplicon.

2. Perform the PCR with your previously designed primers. Include a negative control with dH_2_O instead of DNA.

#### Q5 High-Fidelity PCR reaction master mix

| Reagent | Volume (one sample) |
| --- | --- |
| 5X Q5 Reaction Buffer | 5 µl |
| 10 mM dNTPs | 0.5 µl |
| 10 µM Primer F | 1.25 µl |
| 10 µM Primer R | 1.25 µl |
| dH<sub>2</sub>O | 15.75 µl |
| Q5 High-Fidelity Polymerase | 0.25 µl |
| Total volume | 24 µl (add 1 µl of genomic healthy control DNA) |

#### PCR cycling conditions

| Steps | Temperature | Time | Cycles |
| --- | --- | --- | --- |
| Initial Denaturation | 98°C | 30 s | 1 |
| Denaturation | 98°C | 10 s | 35 cycles |
| Annealing | $T_m$ | 20 s | |
| Extension | 72°C | 25 s/kb of amplicon |  |
| Final extension | 72°C | 2 min | 1 |
| Hold | 10°C | forever |  |

3. While the PCR is running, prepare a 1.0-1.5% agarose gel.

4. Load the 1.0-1.5% agarose gel with 5 µl of PCR product mixed with 1 µl of 6X Loading Dye. Run at 120V for at least 30 minutes.

5. If the band on the agarose gel shows the correct amplicon size with a strong intensity, repeat the PCR with a larger set-up. We usually perform a four times set-up (repeating exactly as above for four identical reactions). Retain and combine the PCR volume of each reaction together in a 1.5 mL Eppendorf (Eppi) tube for Step 6. Refer to our troubleshooting for this step, if multiple bands occur, see Troubleshooting.

6. Clean-up PCR products with a standard PCR clean-up kit. For example, we use GenElute PCR Clean-Up Kit (Sigma Aldrich) according to the manufacturer’s instruction.

**Note:** We recommend eluting with 50 µl of dH_2_O water instead of the elution buffer of the supplied kit.

### Digest PCR product and vector with restriction enzymes

#### Timing: 2 h

Digesting the PCR product and vector with restriction enzymes will create sticky ends for ligation of the amplicon into pET01.

7. Determine the optimum buffer and temperature for the restriction enzymes according to the manufacturer’s instructions. The example below assumes the same enzyme-dependent incubation temperature and buffer is suitable for a double digestion.

8. Prepare the digestion mixture in a 1.5 ml Eppi:

#### Digestion mixture

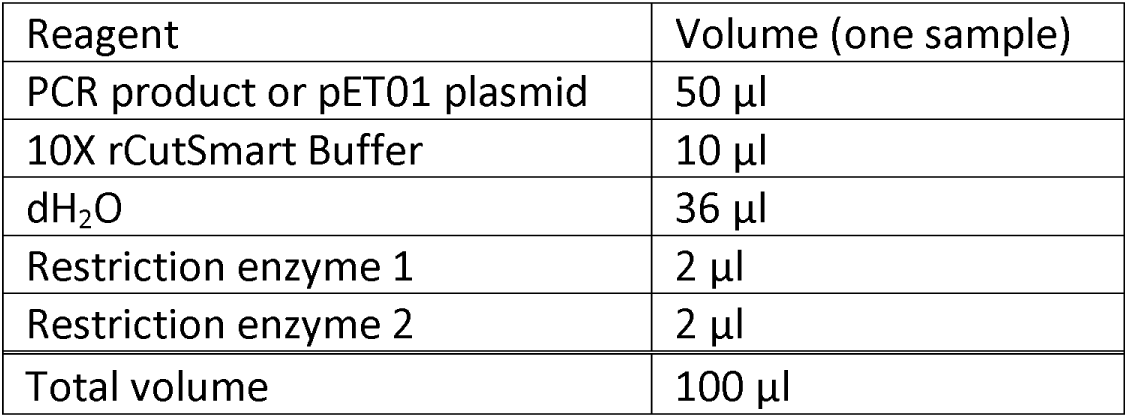

9. Incubate the digestion mixture on a heat block at the enzyme-dependent temperature for 1 hour.

10. Clean-up again with the same kit used in step 6, also eluting with 50 µl of water.

11. Load a 1.0-1.5% agarose gel with 1 µl of the respective digestion mixture, mixed with 5 µl of dH_2_O and 1 µl of 6X Loading Dye. Ensure that the vector is fully digested, as indicated by one band at around 4.2 kb.

**Pause point:** After the clean-up procedure, the PCR product can be stored at -20°C or ligation can be performed immediately after.

**Note:** The digested vector can also be stored at -20°C for later use. As only 1 µl of digested vector is needed for ligation, this can be used for creating more than one pET01 construct.

### Ligate your PCR product into pET01 and perform transformation

#### Timing: 2.5 h + incubation overnight

After ligating the respective insert into the digested pET01 vector, transforming it into competent

*E.coli* will yield a sufficient amount of clones for further analysis prior to transfection.

12. Prepare the ligation mixture in a 1.5 ml Eppi:

#### Ligation mixture

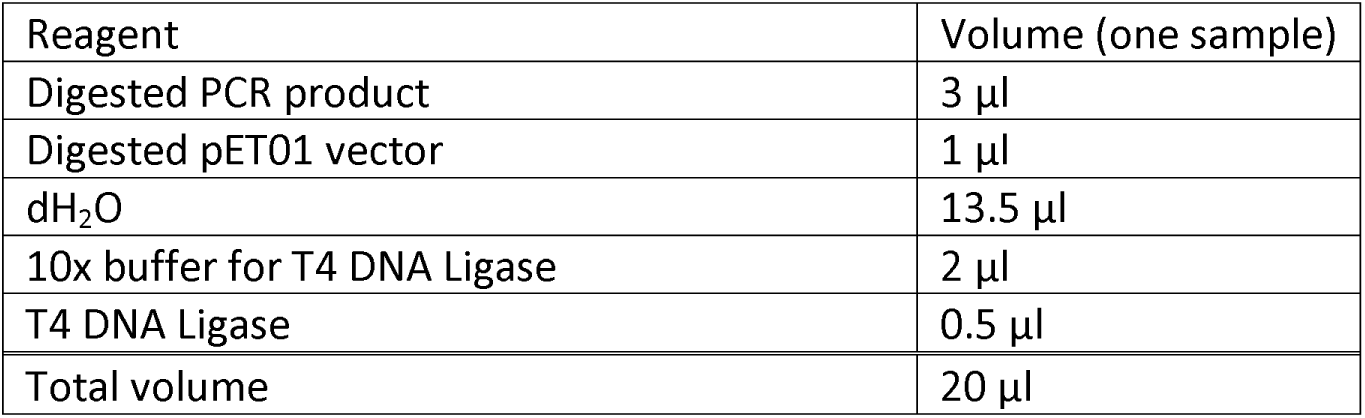

13. Incubate 10 min at room temperature.

**Pause point:** The ligation can also be incubated longer at room temperature, e.g. overnight, or can be stored at -20°C and transformed later following the 10 min incubation.

14. Transform into competent *E.coli* and incubate overnight. We use the following protocol: New England Biolabs: High Efficiency Transformation Protocol (C2987H)^9^.

a. Warm SOC medium to room temperature, put prepared selection plates (LB agar plates with ampicillin) into an incubator to warm to 37°C.
b. Thaw C2987H NEB 5-alpha competent cells on ice for 10 min.
c. Add 5 µl of the ligation mixture onto the cells and carefully flick the tube four to five times to mix. **Critical:** Do not vortex.
d. Incubate on ice for 30 min. Preheat a heat block to 42°C.
e. Heat shock on the heat block for exactly 30 s.
f. Place on ice for 5 min. Adjust heat block to 37°C.
g. Pipette 950 µl of room temperature SOC medium onto the cells.
h. Place the sample onto the heat block for 1 h at 37°C, while shaking (250-300 rpm).
i. Mix the cells thoroughly by flicking the tube and inverting.
j. Dilute the cells in SOC medium. A ∼1:10 dilution usually works (225 µl of SOC medium and 25 µl of cells). **Note:** Keep the rest of the cells stored at room temperature, in case no colonies are observed the next day. Then plate the cells out undiluted. If colonies are observed, freeze the cells with 1:1 40 % glycerol at -80°C as a backup glycerol stock.
k. Spread the diluted cells onto the LB agar selection plates (with ampicillin) and place in incubator at 37°C for 16-24 h.

**Pause point:** The next day, either proceed with the following steps or seal the LB agar plate with parafilm to avoid dehydration, storing it at 4°C for future processing.

### Colony PCR to identify a pET01 clone with insert

#### Timing: 2-4 h

On the LB agar plates containing ampicillin, only *E.coli* clones with the transformed vector should grow. Using a colony screening method, we select a clone that contains the vector with the insert and confirm its identity via PCR, assessing the correct insert size. Subsequent Sanger sequencing will validate this clone.

15. Label a pre-warmed (37°C) new LB agar plate with ampicillin with a grid (called grid plate) and sample name. Calculate the PCR extension time using the formula: 40 s*(0.430 kb + the kilobase pair length of your insert) = extension time in seconds.

16. Calculate the annealing temperature of the primers using the Promega T_m_ Calculator (https://www.promega.de/resources/tools/biomath/tm-calculator/) setting the primer concentration to 300 nM. Use the average of the two calculated temperatures.

17. Prepare a Master Mix depending on the number of colonies you intend to screen. We usually screen five to ten individual colonies. Include a negative control with no colony material added.

#### Colony screen Master Mix

| Reagent | Volume (one sample) |
| --- | --- |
| 5X Reaction Buffer, coloured | 2.5 µl |
| MgCl<sub>2</sub> (50mM) | 0.5 µl |
| dNTPs (100mM) | 0.125 µl |
| 10 µM Primer 02 F | 0.375 µl |
| 10 µM Self-designed insert primer (reverse) | 0.375 µl |
| dH<sub>2</sub>O | 8 µl |
| MangoTaq DNA polymerase | 0.125 µl |
| Total volume | 12 µl |

18. Using a pipette tip, pick one colony from the plate and place the tip onto a square of your grid plate. Insert this tip into a PCR tube containing the Master Mix. Incubate the tip for at least one minute, then re-attach the pipette and pipette up and down to ensure thorough mixing before discarding the tip. Repeat this process for the desired number of colonies to be screened.

19. Incubate the grid plate at 37°C overnight.

20. Run the PCR using the following cycling conditions. Concurrently, prepare a 1.0-1.5% agarose gel.

#### PCR cycling conditions

| Steps | Temperature | Time | Cycles |
| --- | --- | --- | --- |
| Initial Denaturation | 95°C | 5 min | 1 |
| Denaturation | 95°C | 30 s | 35 cycles |
| Annealing | $T_m$ | 30 s | |
| Extension | 72°C | 40 s/kb of amplicon |  |
| Final extension | 72°C | 5 min | 1 |
| Hold | 10°C | forever |  |

21. Load 5 µl of the PCR product mixed with 1 µl of 6X Loading Dye onto the 1.0-1.5% agarose gel. **Critical:** Since the forward primer anneals to vector-specific sequence and the reverse primer anneals to insert-specific sequence, a band should only appear on the agarose gel for colonies containing the vector with insert. The expected band size will be the insert length plus approximately 430 bp, depending on the restriction enzymes used. Absence of a band indicates that the colony does not contain the insert.

22. Select colonies for Sanger sequencing based on gel results, typically choosing three colonies that display a band at the expected size.

**Note**: To conserve resources, consider sequencing the remaining PCR product with standard overnight Sanger sequencing, followed by preparing a single overnight culture after receiving and analyzing the sequencing results. For expedited processing, you may prepare overnight (LB-amp) cultures for all colonies showing the correct band, wait for sequencing the next day and perform plasmid isolation only on the Sanger-confirmed overnight culture.

23. Prepare the samples for sequencing with a standard PCR clean-up reaction. The following shows an example of how this is done but should be adjusted to Sanger sequencing sample requirements in each lab.

#### PCR clean-up Master Mix

| Reagent | Volume (one sample) |
| --- | --- |
| dH<sub>2</sub>O | 3.625 µl |
| Exonuclease I | 0.075 µl |
| FastAP | 0.3 |
| Total volume | 4 µl |

24. Mix 6 µl of the colony screen PCR product with 4 µl of the PCR clean-up Master Mix.

25. Run the following PCR program:

#### PCR clean-up cycling conditions

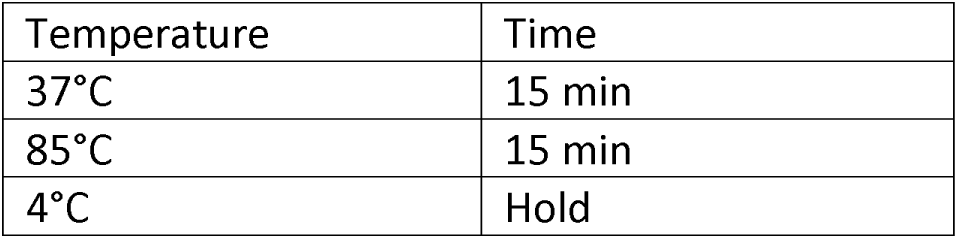

| Temperature | Time |
| --- | --- |
| 37°C | 15 min |
| 85°C | 15 min |
| 4°C | Hold |

26. Prepare a Master Mix for sequencing, which includes the primer for sequencing.

#### Sanger sequencing Master Mix

| Reagent | Volume (one sample) |
| --- | --- |
| dH<sub>2</sub>O | 9 µl |
| PCR primer 02 F | 3 µl |
| Total volume | 12 µl |

**Note:** We usually just use the vector primer PCR primer 02 F for sequencing. If the insert is bigger than 600 base pairs, we also prepare another sample to sequence with the reverse vector primer, which binds to Exon B (PCR primer 03 R).

27. Add 3 µl of the PCR clean up product with one sample of the Sanger sequencing Master Mix into tubes appropriate for your sequencing facility.

28. Confirm the correct insert with analysis of Sanger sequencing results.

### Overnight culture and plasmid isolation

#### Timing: 10 min + 12-18 h incubation + 1.5 h plasmid isolation

After confirming a clone with the correct insert with Sanger sequencing, set up an overnight culture.

**Critical:** Compared to other vectors used in our lab, we have observed lower pET01 vector concentration yields when using standard Miniprep kits for plasmid isolation. To avoid the need for repeated plasmid isolations, particularly in cases where repeat transfections may be needed, duplicating/triplicating transfections or transfecting the same construct in multiple cell lines, we advise using a Midiprep kit (e.g. GenElute Plasmid Midiprep Kit (Sigma-Aldrich)) for higher yield.

29. Aliquot 20 ml of LB medium into a suitable overnight tube and add 10.5 µl of ampicillin (concentration=50 mg/ml).

30. Using a pipette tip, inoculate the Sanger confirmed colony from the grid plate into the LB medium with ampicillin.

**Pause point:** Incubate the culture for 14-18 hours at 37°C, while shaking at 150-400 rpm in an incubator.

**Note:** After incubation freeze 500 µl of the overnight culture with 500 µl of 40 % glycerol in a vial suitable for freezing at -80°C for back-up. Process the remaining sample in Step 31.

31. Follow the manufacturer’s instructions of the Midiprep kit for plasmid isolation with following modification:

a. For a more concentrated eluate, use 500 µl of preheated (65°C) dH_2_O as the elution solution and incubate on the column for 10 min before elution.

32. Measure the concentration of your isolated plasmid using a Nanodrop spectrophotometer.

### Create a variant minigene via Site-Directed Mutagenesis

#### Timing: 3-4 days

After isolating the wild-type construct, the following PCR-based mutagenesis method allows for simple introduction of specific variants using pre-designed mutagenesis primers. This technique can facilitate insertions, deletions or single-base pair exchanges. We follow the protocol available at dx.doi.org/10.17504/protocols.io.bddfi23n^10^.

33. If not already previously done, measure the concentration of the wild-type template plasmid. Dilute the plasmid to a concentration of 20 ng/µl using dH_2_O in a total volume of 15 µl.

34. Prepare the following Master Mix with standard components supplied in the Q5 Site-Directed Mutagenesis Kit:

#### Q5 SDM Master Mix

| Reagent | Volume (one sample) |
| --- | --- |
| Q5 Hot Start High-Fidelity 2X Master Mix | 12.5 µl |
| 10 µM Mutagenesis Primer (forward) | 1.25 µl |
| 10 µM Mutagenesis Primer (reverse) | 1.25 µl |
| Wild-type plasmid (20 ng/µl) | 1 µl |
| dH<sub>2</sub>O | 9 µl |
| Total volume | 25 µl |

35. Mix by pipetting up and down and spin down shortly in a table centrifuge.

36. Calculate the annealing temperature and the extension time of your PCR. We use the annealing temperature provided by the NEBaseChanger website (https://nebasechanger.neb.com/).

37. Run the following PCR program according to the manufacturer protocol:

#### PCR cycling conditions

| Steps | Temperature | Time | Cycles |
| --- | --- | --- | --- |
| Initial Denaturation | 98°C | 30 s | 1 |
| Denaturation | 98°C | 10 s | 25 cycles |
| Annealing | T<sub>m</sub> | 10 s |  |
| Extension | 72°C | 20 s/kb of plasmid |  |
| Final extension | 72°C | 2 min | 1 |
| Hold | 4-10°C | forever |  |

38. Prepare for a transformation by performing Step 14a. Thaw a tube of NEB 5-alpha competent *E. coli* on ice for 5 minutes. Assemble the following reagents in a 1.5 ml Eppendorf tube, then gently pipette up and down.

#### KLD enzyme Mix

| Reagent | Volume (one sample) |
| --- | --- |
| PCR Product | 1 µl |
| 2X Reaction Buffer | 5 µl |
| 10X KLD Enzyme Mix | 1 µl |
| dH<sub>2</sub>O | 3 µl |
| Total volume | 10 µl |

39. Incubate the KLD enzyme reaction mixture at room temperature for 5 min, while allowing *E. coli* cells to thaw on ice for the same period (10 minutes total incubation time).

40. Perform transformation starting with step 14c by adding 5 µl of the KLD enzyme mixture. **Pause point:** Incubate the plate overnight at 37°C. The next day, either proceed with the following steps or seal the LB agar plate with parafilm to avoid dehydration, storing it at 4°C for future processing.

41. Screen the colonies for the desired insert, following the same procedure from steps 15-28.

42. After validating a clone containing the correct insert and variant, set-up an overnight culture for plasmid isolation the next day (same as steps 29-32).

### Transfecting minigenes

#### Timing: 30 min (day 1) + 30 min (day 2)

Transfect wild-type and variant plasmid constructs into the desired cell line to utilize the inherent transcription and splicing processes of the cells. Optimize the transfection procedure for your cell line of choice. This protocol describes transient transfection in HCT116 cells with RPMI medium. Results were replicated in HEK293T cells, using DMEM medium instead.

**Critical:** Seed 4×10^5^ cells per well in a 6-well plate the day before transfection, ensuring 60-70 % confluence on the day of transfection.

**Note:** For transfection, include not only the wild-type and variant construct, but also an empty pET01 vector control and a transfection negative control.

43. Prewarm FBS-free RPMI medium to 37°C.

44. In a 1.5 ml Eppi, dilute 1 µg of each isolated plasmid construct with FBS-free RPMI medium to a total volume of 95 µl.

45. Add 3 µl of FuGene 6 Transfection Reagent to each transfection mix. Gently mix and incubate at room temperature for 15 min.

46. During incubation, change the medium in the 6-well plate to 2ml of pre-warmed FBS-free RPMI medium per well.

47. Slowly add the transfection mix dropwise to the respective well using a pipette. Gently mix the plate by tilting and incubate at 37°C humidified incubator with 95% humidity and 5% CO_2_ for 24 h.

48. 24 hours post-transfection change the medium back to supplemented medium (with FBS). Incubate under the same conditions for another 24 h.

### Isolate RNA and synthesize cDNA

#### Timing: 3-4 h + 2.5 h incubation

48 hours post-transfection, isolate RNA from your transfected cells two-step RT-PCR. For the first step use a primer with sequence provided by the manufacturer that anneals to the pET01 vector sequence (step 52). For the second step, use primers amplifying across exons A and B (step 56).

49. Isolate total RNA 48h post-transfection using a standard RNA isolation kit. We recommend Qiagen’s miRNeasy Mini Kit and protocol with the following modifications:

**Critical**: Pre-chill centrifuge to 4°C and pre-warm RNase free water to 55°C.

a. Remove the medium from the well plate and add 2 ml of DPBS in each well.
b. Detach the cells by pipetting and transfer to 2 ml Eppis.
c. Pellet the cells by centrifugation at 3,000 g and 4°C for 5 min.
d. Discard the supernatant and flick the Eppi to dissolve the pellet in the residual DPBS.
e. Add 700 µl of QIAzol Lysis Reagent and vortex for 1 min.
f. Incubate at room temperature for 5 min.
g. Add 140 µl of chloroform and shake vigorously for 15 s.
h. Incubate for 5 min at room temperature.
i. Centrifuge for 15 min at 12,000 g at 4°C. **Critical:** Ensure the formation of three phases: A pink organic phase (bottom), a white interphase and an aqueous top phase containing the RNA. If the phases are unclear, re-centrifuge or check if all components were added.
j. Transfer 350 µl of the upper phase to a new Eppi and mix with 525 µl 100% ethanol. **Critical:** Do not transfer the white interphase or pink organic phase. If this proves difficult, transfer a smaller volume of the upper phase or re-centrifuge ensure separation of the phases. **Note:** From now on warm the centrifuge to room temperature again or use another centrifuge at room temperature.
k. Transfer 700 µl of the mixture to the provided column and centrifuge for 15 s at 10,000 g at room temperature.
l. Decant the flow-through.
m. Add the remaining mixture from step j to the column and centrifuge for 15 s at 10,000 g (room temperature) and decant the flow-through.
n. Add 700 µl of RWT buffer to the column. Centrifuge for 15 s at 10,000 g (room temperature) and decant the flow-through.
o. . Add 500 µl of the RPE buffer to the column. Centrifuge for 15 s at 10,000 g (room temperature) and decant the flow-through.
p. Repeat step o.
q. Transfer the column to a new collection tube and centrifuge for 1 min at full speed (room temperature).
r. Transfer the column to a new Eppi and add 35 µl of the pre-warmed RNase free water directly onto the center of the column.
s. Incubate for 1 min.
t. Centrifuge for 1 min at 10,000 g (room temperature) to elute the RNA.

**Critical**: Always keep the isolated RNA on ice from this point forward.

50. Measure the RNA concentration with a Nanodrop spectrophotometer, ensuring RNase free water is used as the blank. See Troubleshooting

**Pause point:** Store the RNA at -80°C or proceed immediately with step 51. When continuing directly with step 51, store any remaining RNA at -80°C.

51. Dilute the RNA in RNase free water to 2 µg in a volume of 10 µl.

52. Prepare a RT-PCR Master Mix with the High-Capacity cDNA Reverse Transcription Kit (Applied Biosystems) and the cDNA primer 01 on ice:

#### RT-PCR Master Mix

| Reagent | Volume (one sample) |
| --- | --- |
| 10x RT Buffer | 2 µl |
| 25x dNTP Mix (100mM) | 0.8 µl |
| cDNA Primer 01 (100 µM) | 2 µl |
| MultiScribe Reverse Transcriptase | 1.0 µl |
| RNase free H<sub>2</sub>O | 4.2 µl |
| Total volume | 10 µl |

53. Gently mix and aliquot in individual PCR tubes.

54. Pipette 10 µl of RNA (2 µg) into the PCR tube, gently mix by pipetting.

55. Run the PCR with following program:

#### PCR cycling conditions

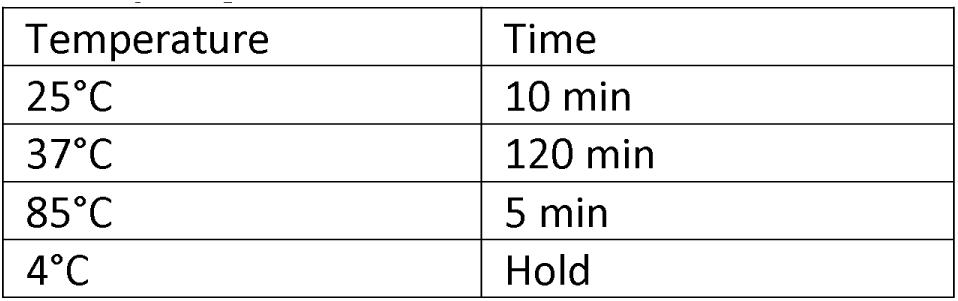

| Temperature | Time |
| --- | --- |
| 25°C | 10 min |
| 37°C | 120 min |
| 85°C | 5 min |
| 4°C | Hold |

**Pause point**: Store labeled cDNA at -20°C until ready to perform the final PCR with PCR primer 02 F and 03 R. This will amplify the region between Exon A and Exon B of the pET01 vector.

56. Prepare a Master Mix for the final PCR with a standard PCR kit.

**Critical:** If fragment analysis is required to quantify different amplicons, use one FAM-labeled primer for this PCR (e.g. FAM-labeled forward primer and a non-FAM-labeled reverse primer). Additionally, to prevent interference of FAM-labeled amplicons with sequencing, run a parallel PCR with non-FAM-labeled primers. Be sure to include controls.

**Note:** We obtained good PCR results using the QIAGEN Multiplex PCR Kit. However, depending on the sample other PCR kits might be more suitable. Refer to our troubleshooting guide if problems occur, see Troubleshooting.

#### Final PCR Master Mix

| Reagent | Volume (one sample) |
| --- | --- |
| 2x QIAGEN Multiplex PCR Master Mix | 5 µl |
| 10 µM (FAM-)PCR primer 02 F | 1 µl |
| 10 µM PCR primer 03 R | 1 µl |
| dH<sub>2</sub>O | 11.5 µl |
| Add cDNA | 1.5 µl |
| Total volume | 20 µl |

57. Run a touchdown PCR program. Make adjustments if needed.

#### PCR cycling conditions

| Step | Temperature | Time | Cycles |
| --- | --- | --- | --- |
| 1 | 95°C | 5 min. | 1 |
| 2 | 95°C | 30 s | 1 |
| 3 | 62°C | 30 s | 1 |
| 4 | 72°C | 30 s | Go to step 2, repeat 3 times |
| 5 | 95°C | 30 s | 1 |
| 6 | 59°C | 30 s | 1 |
| 7 | 72°C | 30 s | Go to step 5, repeat 5 times |
| 8 | 95°C | 30 s | 1 |
| 9 | 56°C | 30 s | 1 |
| 10 | 72°C | 30 s | Go to step 8, repeat 30 times |
| 11 | 72°C | 8 min | 1 |
| 12 | 10°C | hold |  |

58. Prepare a 1.0-1.5% agarose gel and run the PCR products with 6X Loading Dye for at least 1 h at 120V. See expected outcomes for an example gel.

### Optional: Cloning to isolate single sequences in instances where splicing yields multiple amplicons

#### Timing: variable

If multiple bands are observed on the final agarose gel, direct sequencing of the RT-PCR product may result in noisy or unusable sequencing results. To avoid this, you can either clone the RT-PCR products into another vector or perform gel extraction.

**Note**: We have achieved good results with the QIAquick Gel Extraction Kit (QIAGEN). However, if the splice products differ by only a very small number of base pairs, cloning the final PCR products into a vector is preferable for cleaner sequencing results. For instructions on subsequent cloning of the RT-PCR products into the pCR™4Blunt TOPO™ vector (Fisher Scientific), follow the next steps.

**Critical:** Ensure that the RT-PCR products used in these steps are generated with non-FAM labeled primers. Use 30-60 µl of PCR product (depending on its intensity on the agarose gel) and purify it using a standard PCR clean-up column kit. Elute the product in 30 µl of dH_2_0.

59. Ligate the RT-PCR product into the pCR™4Blunt TOPO™ vector:

#### Ligation mixture

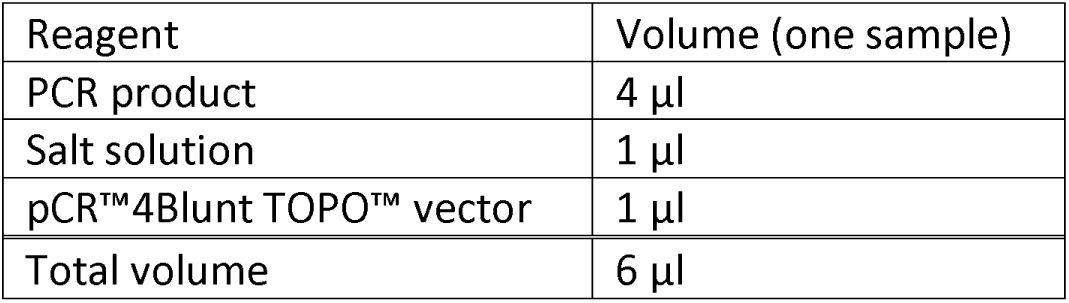

| Reagent | Volume (one sample) |
| --- | --- |
| PCR product | 4 µl |
| Salt solution | 1 µl |
| pCR™4Blunt TOPO™ vector | 1 µl |
| Total volume | 6 µl |

60. Gently mix the ligation reaction, briefly spin down in a tabletop centrifuge and incubate for 15-30 min at room temperature.

61. Transform all 6 µl into NEB 5-alpha Competent *E.coli* described in step 14. This time, plate out 250 µl of undiluted transformed bacteria onto LB agar plates (with kanamycin).

**Pause point:** Incubate 24 h at 37°C.

62. Perform a colony PCR and send the PCR products for Sanger sequencing, as described in steps 16-28. This time use M13 forward and reverse primers.

### Quantification

Fragment Analysis is used to quantify different sizes of PCR products labeled with fluorescent dye (FAM label) using capillary electrophoresis. It is especially useful to detect small quantities of splice products that cannot be visualized on an agarose gel. Furthermore, fragment analysis by capillary array electrophoresis achieves high sensitivity to allow for relative quantification of multiple bands and it is especially helpful for providing high resolution to detect base pair differences of 1 bp or more that may not be easily resolved on agarose gels.

63. Prepare FAM-labeled final PCR products as in steps 56-58 for Fragment Analysis.

64. Dilute samples with suitable buffer according to internal lab standard.

65. Mix samples with “orange dye” labeled size standard according to internal lab standard.

66. Denature samples at 96°C for 4 min.

67. Load samples onto the fragment analyzer. We used the ABI3730 XL DNA analyzer with the following conditions: injection voltage of 1.6 V and injection time of 15 s, run voltage of 15 kV and run time of 2500 s.

68. Analyze data with fragment analysis software such as GeneMapper 5 (Applied Biosystems).

## Expected outcomes

In this protocol, we used the pET01 vector to study the splicing effects of inherited patient variants in *OTOF*. Variants can affect pre-mRNA splicing through various mechanisms, leading to different consequences. Disruptions to the highly conserved 5’ splice donor, 3’ splice acceptor sites or splicing regulatory elements (SREs) can interfere with the recognition by trans-acting splicing factors.

Additionally, some variants can also activate cryptic splice sites. The consequences of these disruptions can include exon skipping, partial or complete intron retention or exon truncation. These events can result in insertions or deletions, which may either preserve or disrupt the reading frame, or in frameshifts that introduce a premature stop codon, typically resulting in a truncated and often non-functional protein.^1,11^

When analyzing splicing patterns in the minigene model, it is essential to compare results to the wild-type minigene, which should express the correctly spliced exons. Figure 2 illustrates exon skipping caused by a variant in *OTOF.* Examples of variants assayed with this protocol and results shown in two cell lines are accessible in the supplementary material (Figure S2-S5).

**Figure 2:**
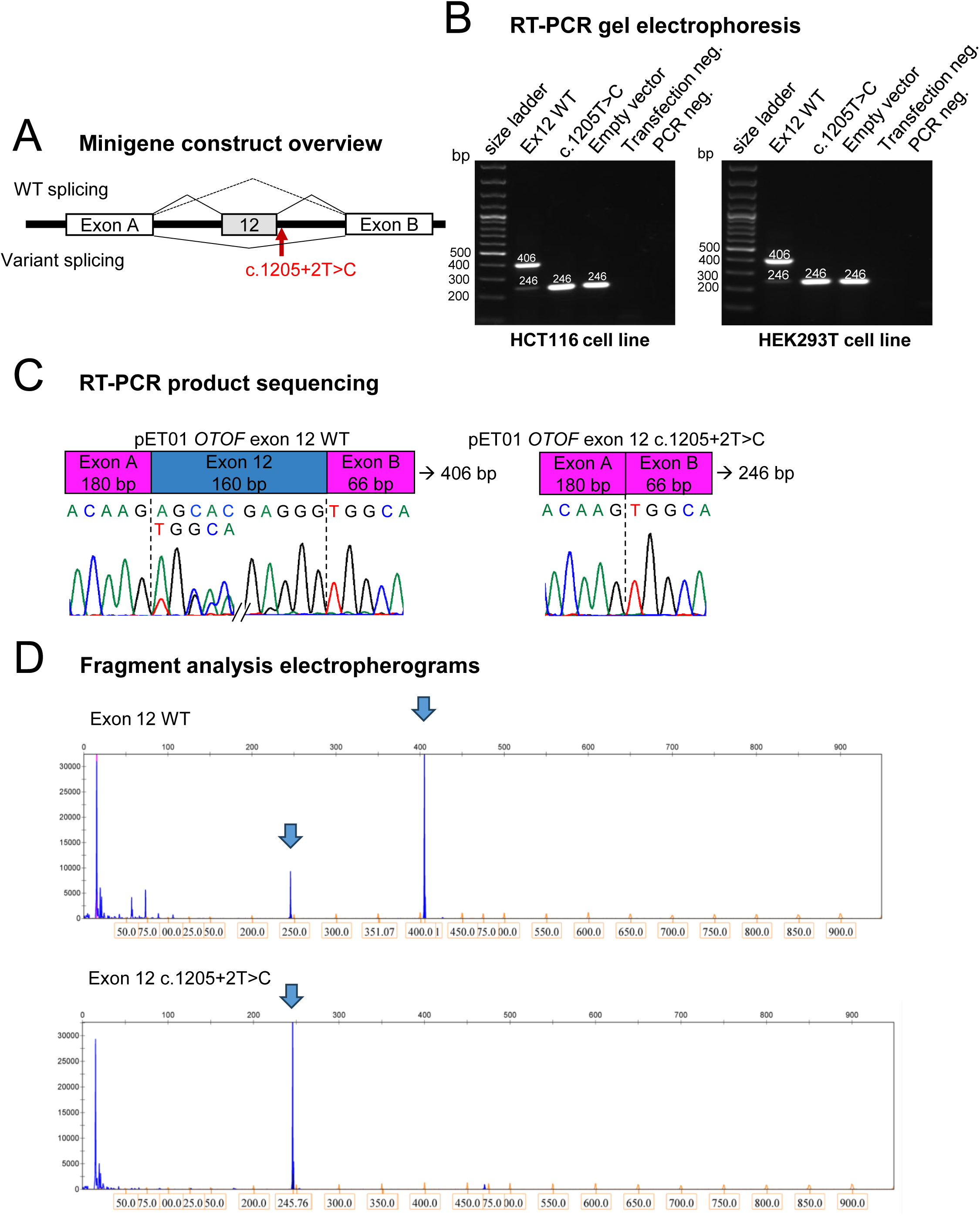
Minigene assay of the c.1205+2T>C *OTOF* variant. (A) Minigene construct schematic with Exon 12 of *OTOF* cloned in between exon A and B of the vector. The red arrow displays the position of the variant. The WT and variant construct splicing pattern is indicated on top and bottom, respectively. (B) Visualization of final RT-PCR products after gel electrophoresis. Minigenes were transfected into HCT116 (left) and HEK293T (right) cells. (C) Direct Sanger sequencing results of RT-PCR products of HCT116 cells. The primer PCR 02 F, aligning to exon A of pET01 was used as a primer for sequencing. Exon A and B of the vector are indicated by pink boxes, exon 12 by a blue box. The wild-type construct sample shows normal exon 12 splicing and skipping of exon 12. The variant sample shows exon 12 skipping only. (D) Electropherogram of one triplicate of the final RT-PCR products from HCT116 cells. The blue arrows point to peaks that correspond to the sizes indicated on the agarose gel in panel B.

## Limitations

While the minigene assay is a widely used, robust, quick and cost-effective method for assessing splice effects of genetic variants and it provides validation of splice effects on the RNA-level, it has certain limitations.^12,13^

The insert for the minigene is only a small fragment of the full-length gene, which can be an advantage to reduce complexity and focus on the most important regions. However, this simplification can also be a drawback, as the absence of the full genomic context may cause regulatory elements located far from the splice sites to be missed, which can influence splicing. Another potential limitation is the choice of cell line for transfection, as it may affect splicing outcomes. Ideally, if available, a cell line that closely mimics the tissue-specific splicing of the gene of interest should be used to potentially improve the relevance of the results.

In addition, this assay requires cloning, transfection and subsequent analysis, which can be labour and resource intensive.

Furthermore, while analysis of mRNA in this assay provides valuable insight into changes in splice patterns, interpreting the subsequent protein presence and function should be evaluated with caution due to many factors influencing translation and post-translational modifications.

Some mis-splicing events, such as exon skipping, have been observed even when expressing the wild-type minigene, likely due to the loss of regulatory sequences within the intronic regions included in the construct or the strong artificial splice sites of the vector. This limitation, combined with the size restrictions of the cloned DNA segments, makes minigene assays potentially unsuitable for functionally assessing deep intronic variants in very large introns. Therefore, each variant requires careful evaluation to determine the most appropriate functional assay to use.

While our experimental findings highlight some limitations in minigene-based splicing analysis, we have validated the use of a customizable minigene assay within clinical investigations and successfully identified variants causing aberrant splicing. In the absence of appropriate patient samples for RNA investigation, this is a critical tool for clinical variant investigation.

## Troubleshooting

**Problem 1:**

The first PCR product presents as multiple bands on the agarose gel (major step 1).

**Potential solution:**

- Perform a gradient PCR with increased T_m_.
- Choose a different DNA sample (if available) to avoid SNPs.
- Design new primers.

**Problem 2:**

The final RT-PCR product shows no band on the agarose gel (step 56-58).

**Potential solution:**

Check cDNA quality and repeat from major step 8 forward.

Consider using a different PCR kit and optimizing PCR conditions. If the wild-type exon is not expressed, consider re-designing the assay to include a larger intronic region.

**Problem 3:**

The electropherogram in fragment analysis always shows double peaks.

**Potential solution:**

Make sure to use the final PCR product with only one FAM-labeled primer.

**Problem 4:**

The RNA concentration is too low at the end of the RNA extraction.

**Potential solution:**

Always optimize transfection conditions for the cell line of use, e.g. by increasing or decreasing incubation time after transfection or by increasing or decreasing the concentration of transfected plasmid.

## Resource availability

### Lead contact

Further information and requests for resources should be directed to the lead contact, Barbara Vona.

### Technical contact

Questions about the technical specifics of performing the protocol should be directed to and will be answered by the technical contact, Hannah Andreae.

### Materials availability

Further requests for resources should be directed to the lead contact, Barbara Vona.

### Data and code availability

This study did not generate code. Sequencing data and a dataset generated with this protocol is included.

## Acknowledgments

We are thankful to the German Research Foundation DFG VO 2138/7-1 grant 469177153 that supported this work. We thank Dayana Warnecke at the Department of Molecular Neurobiology, Max Planck Institute for Multidisciplinary Sciences for excellent technical support. BV is a member of the European Reference Network on Rare Congenital Malformations and Rare Intellectual Disability ERN-ITHACA [EU Framework Partnership Agreement ID: 3HP-HP-FPA ERN-01-2016/739516]. This work was done with support of the Center for Rare Hearing Disorders at the Center of Rare Diseases Göttingen (ZSEG).

## Author contributions

HA and FB performed the experiments. HA wrote the first draft. All authors participated in review and editing. BV conceived the project and designed the experiment. All authors have read and agreed to the content.

## References

1. Cartegni, L., Chew, S.L., and Krainer, A.R. (2002). Listening to silence and understanding nonsense: exonic mutations that affect splicing. Nat Rev Genet 3, 285–298. 10.1038/nrg775.

2. Truty, R., Ouyang, K., Rojahn, S., Garcia, S., Colavin, A., Hamlington, B., Freivogel, M., Nussbaum, R.L., Nykamp, K., and Aradhya, S. (2021). Spectrum of splicing variants in disease genes and the ability of RNA analysis to reduce uncertainty in clinical interpretation. Am J Hum Genet 108, 696–708. 10.1016/j.ajhg.2021.03.006.

3. Yasunaga, S., Grati, M., Chardenoux, S., Smith, T.N., Friedman, T.B., Lalwani, A.K., Wilcox, E.R., and Petit, C. (2000). OTOF Encodes Multiple Long and Short Isoforms: Genetic Evidence That the Long Ones Underlie Recessive Deafness DFNB9. Am J Hum Genet 67, 591–600.

4. MobiTec GmbH Exontrap.

5. Davis, M.W., and Jorgensen, E.M. (2022). ApE, A Plasmid Editor: A Freely Available DNA Manipulation and Visualization Program. Front Bioinform 2, 818619. 10.3389/fbinf.2022.818619.

6. Harrison, P.W., Amode, M.R., Austine-Orimoloye, O., Azov, A.G., Barba, M., Barnes, I., Becker, A., Bennett, R., Berry, A., Bhai, J., et al. (2024). Ensembl 2024. Nucleic Acids Research 52, D891–D899. 10.1093/nar/gkad1049.

7. Raney, B.J., Barber, G.P., Benet-Pagès, A., Casper, J., Clawson, H., Cline, M.S., Diekhans, M., Fischer, C., Navarro Gonzalez, J., Hickey, G., et al. (2024). The UCSC Genome Browser database: 2024 update. Nucleic Acids Research 52, D1082–D1088. 10.1093/nar/gkad987.

8. Untergasser, A., Cutcutache, I., Koressaar, T., Ye, J., Faircloth, B.C., Remm, M., and Rozen, S.G. (2012). Primer3—new capabilities and interfaces. Nucleic Acids Research 40, e115–e115. 10.1093/nar/gks596.

9. England Biolabs, N. (2020). High Efficiency Transformation Protocol (C2987H) v3. Preprint, 10.17504/protocols.io.bddti26n 10.17504/protocols.io.bddti26n.

10. England Biolabs, N. (2020). Q5® Site-Directed Mutagenesis (E0554) v2. Preprint, 10.17504/protocols.io.bddfi23n 10.17504/protocols.io.bddfi23n.

11. Abramowicz, A., and Gos, M. (2018). Splicing mutations in human genetic disorders: examples, detection, and confirmation. J Appl Genet 59, 253–268. 10.1007/s13353-018-0444-7.

12. Sharma, N., Sosnay, P.R., Ramalho, A.S., Douville, C., Franca, A., Gottschalk, L.B., Park, J., Lee, M., Vecchio-Pagan, B., Raraigh, K.S., et al. (2014). Experimental assessment of splicing variants using expression minigenes and comparison with in silico predictions. Hum Mutat 35, 1249–1259. 10.1002/humu.22624.

13. Martin, N., Bergougnoux, A., Baatallah, N., Chevalier, B., Varilh, J., Baux, D., Costes, B., Fanen, P., Raynal, C., Sermet-Gaudelus, I., et al. (2021). Exon identity influences splicing induced by exonic variants and in silico prediction efficacy. J Cyst Fibros 20, 464–472. 10.1016/j.jcf.2020.12.003.

