## Supplementary material for "Protocol for a minigene splice assay using the pET01 vector": Figure S1

### Exontrap vector pET01-full plasmid sequencing

Exon A and B of pET01
pET01 PCR primer 02 F: with BamHI restriction site
pET01 PCR primer 03 R:  with SmaI restriction site
cDNA primer 01
XhoI restriction site
BamHI restriction site
SalI restriction site
NotI restriction site
ApaI restriction site
SmaI restriction site
SpeI restriction site
XbaI restriction site
SacII restriction site

AACGCCAGCAACGGAGATGCGCCGCGTGCGGCTGCTGGAGATGGCGGACGCGATGGATATGTTCTGCC
AAGGGTTGGTTTGCGCATTCACAGTTCTCCGCAAGAATTGATTGGCTCCAATTCTTGGAGTGGTGAATCC
GTTAGCGAGGTGCCGCCGGCTTCCATTCAGGTCGAGGTGGCCCGGCTCCATGCACCGCGACGCAACGCG
GGGAGGCAGACAAGGTATAGGGCGGCGCCTACAATCCATGCCAACCCGTTCCATGTGCTCGCCGAGGC
GGCATAAATCGCCGTGACGATCAGCGGTCCAATGATCGAAGTTAGGCTGGTAAGAGCCGCGAGCGATC
CTTGAAGCTGTCCCTGATGGTCGTCATCTACCTGCCTGGACAGCATGGCCTGCAACGCGGGCATCCCGAT
GCCGCCGGAAGCGAGAAGAATCATAATGGGGAAGGCCATCCAGCCTCGCGTCGGGGAGCTTTTTGCAA
AAGCCTAGGCCTCCAAAAAAGCCTCCTCACTACTTCTGGAATAGCTCAGAGGCCGAGGCGGCCTCGGCC
TCTGCATAAATAAAAAAAATTAGTCAGCCATGGGGCGGAGAATGGGCGGAACTGGGCGGAGTTAGGGG
CGGGATGGGCGGAGTTAGGGGCGGGACTATGGTTGCTGACTAATTGAGATGCATGCAAGGAGATGGC
GCCCAACAGTCCCCCGGCCACGGGGCCTGCCACCATACCCACGCCGAAACAAGCGCTCATGAGCCCGAA
GTGGCGAGCCCGATCTTCCCCATCGGTGATGTCGGCGATATAGGCGCCAGCAACCGCACCTGTGGCGCC
GGTGATGCCGGCCACGATGCGTCCGGCGTAGAGGATCTCAGGATATAGTAGTTTCGCTTTTGCATAGGG
AGGGGGAAATGTAGTCTTATGCAATACTCTTGTAGTCTTGCAACATGCTTATGTAACGATGAGTTAGCAA
CATGCCTTATAAGGAGAGAAAAAGCACCGTGCATGCCGATTGGTGGGAGTAAGGTGGTATGATCGTGG
TATGATCGTGCCTTGTTAGGAAGGCAACAGACGGGTCTAACACGGATTGGACGAACCACTGAATTCCGC
ATTGCAGAGATATTGTATTTAAGTGCCTAGCTCGATACAATAAACGCCATTTGACCATTCACCACATTGGT
GTGCACCTCAAGCTTCCTGCATGCTGCTGCTGCTGCTGCTGGGCCTGAGGCTACAGCTCTCCCTGGG
CATCATCCCAGTTGAGGAGGAGAACCCGGACTTCTGGAACCGCGAGGCAGCCGAGGCCCTGGGTGCCG
CCAAGAAGCTGCAGCCTGCACAGACAGCCGCCAAGAACCTCATCATCTTCCTGGGCGATGGGATGGGGG
TGTCTACGGTGACAGCTGCCAGGATCGATCCGCTTCCTGCCCCTGCTGGCCCTGCTCATCCTCTGGGAGC
CCCGCCCTGCCCAGGCTTTTGTCAAACAGCACCTTTGTGGTTCTCACTTGGTGGAAGCTCTCTACCTGGTG
TGTGGGGAGCGTGGATTCTTCTACACACCCATGTCCCGCCGCGAAGTGGAGGACCCACAAGGTAAGCTC
TGCTCCTGAATTAATTCTATCCCAAGTGCTAACTACCCTGTTTGTCTTTCACCCTTGAGACCTTGTAAATTG
TGCCCTAGGTGTGGAGGGTCTCAGGCTAACCAGTGGGGGGCACATTTCTGTGGGCAGCTAGACATATGT
AAACATGGTAGCTGCCAGGAAGGAGTGAGAATCCTTCCTTAAGTCTCCTAGGTGGTGACGGGTGGCTAG
GCCCCAGGATAGGTACCGGGCCCCCCCTCGAGGTCGACGGTATCGATAAGCTAATTCCTGCAGCCCGGG
GGATCCACTAGTTCTAGAGCGGCCGCCACCGCGGTGGAGCTCGGTACCTATTTGGGGACCCCATAGAGC

ACTGCACTGACTGAGGGATGGTAACAGGATGTGTAGGTTTTGGAGGCCCATATGTCCATTCATGACCAG
TGACTTGTCTCACAGCCATGCAACCCTTGCCTCCTGTGCTGACTTAGCAGGGGATAAAGTGAGAGAAAGC
CTGGGCTAATCAGGGGGTCGCTCAGCTCCTCCTAACTGGATTGTCCTATGTGTCTTTGCTTCTGTGCTGCT
GATGCTCTGCCCTGTGCTGACATGACCTCCCTGGCAGTGGCACAACTGGAGCTGGGTGGAGGCCCGTGA
CCTTCAGACCTTGGCACTGGAGGTGGCCCGGCAGAAGCGCGGCATCGTGGATCAGTGCTGCACCAGCAT
CTGCTCTCTCTACCAACTGGAGAACTACTGCAACTAGGCCCACCACTACCCTGTCCACCCCTCTGCAATGA
ATAAAACCTTTGAAAGAGCACTACAAGTTGTGTGTACATGCGTGCATGTGCATATGTGGTGCGGGGGGA
ACATGAGTGGGGCTGGCTGGAGTGGCGATGATAAGCTGTCAAACATGAGAATTCTTGAAGACGAAAGG
GCCTCGTGATACGCCTATTTTTATAGGTTAATGTCATGATAATAATGGTTTCTTAGACGTCAGGTGGCACT
TTTCGGGGAAATGTGCGCGGAACCCCTATTTGTTTATTTTTCTAAATACATTCAAATATGTATCCGCTCAT
GAGACAATAACCCTGATAAATGCTTCAATAATATTGAAAAAGGAAGAGTATGAGTATTCAACATTTCCGT
GTCGCCCTTATTCCCTTTTTTGCGGCATTTTGCCTTCCTGTTTTTGCTCACCCAGAAACGCTGGTGAAAGTA
AAAGATGCTGAAGATCAGTTGGGTGCACGAGTGGGTTACATCGAACTGGATCTCAACAGCGGTAAGATC
CTTGAGAGTTTTCGCCCCGAAGAACGTTTTCCAATGATGAGCACTTTTAAAGTTCTGCTATGTGGCGCGG
TATTATCCCGTGTTGACGCCGGGCAAGAGCAACTCGGTCGCCGCATACACTATTCTCAGAATGACTTGGT
TGAGTACTCACCAGTCACAGAAAAGCATCTTACGGATGGCATGACAGTAAGAGAATTATGCAGTGCTGC
CATAACCATGAGTGATAACACTGCGGCCAACTTACTTCTGACAACGATCGGAGGACCGAAGGAGCTAAC
CGCTTTTTTGCACAACATGGGGGATCATGTAACTCGCCTTGATCGTTGGGAACCGGAGCTGAATGAAGCC
ATACCAAACGACGAGCGTGACACCACGATGCCTGCAGCAATGGCAACAACGTTGCGCAAACTATTAACT
GGCGAACTACTTACTCTAGCTTCCCGGCAACAATTAATAGACTGGATGGAGGCGGATAAAGTTGCAGGA
CCACTTCTGCGCTCGGCCCTTCCGGCTGGCTGGTTTATTGCTGATAAATCTGGAGCCGGTGAGCGTGGGT
CTCGCGGTATCATTGCAGCACTGGGGCCAGATGGTAAGCCCTCCCGTATCGTAGTTATCTACACGACGGG
GAGTCAGGCAACTATGGATGAACGAAATAGACAGATCGCTGAGATAGGTGCCTCACTGATTAAGCATTG
GTAACTGTCAGACCAAGTTTACTCATATATACTTTAGATTGATTTAAAACTTCATTTTTAATTTAAAAGGAT
CTAGGCTGCTGCTTGCAAACAAAAAAACCACCGCTACCAGCGGTGGTTTGTTTGCCGGATCAAGAGCTAC
CAACTCTTTTTCCGAAGGTAACTGGCTTCAGCAGAGCGCAGATACCAAATACTGTCCTTCTAGTGTAGCC
GTAGTTAGGCCACCACTTCAAGAACTCTGTAGCACCGCCTACATACCTCGCTCTGCTAATCCTGTTACCAG
TGGCTGCTGCCAGTGGCGATAAGTCGTGTCTTACCGGGTTGGACTCAAGACGATAGTTACCGGATAAGG
CGCAGCGGTCGGGCTGAACGGGGGGTTCGTGCACACAGCCCAGCTTGGAGCGAACGACCTACACCGAA
CTGAGATACCTACAGCGTGAGCTATGAGAAAGCGCCACGCTTCCCGAAGGGAGAAAGGCGGACAGGTA
TCCGGTAAGCGGCAGGGTCGGAACAGGAGAGCGCACGAGGGAGCTTCCAGGGGGAAACGCCTGGTAT
CTTTATAGTCCTGTCGGGTTTCGCCACCTCTGACTTGAGCGTCGATTTTTGTGATGCTCGTCAGGGGGGC
GGAGCCTATGGAAA
