## Supplementary material for "Protocol for a minigene splice assay using the pET01 vector": Figure S2

### Slide 1
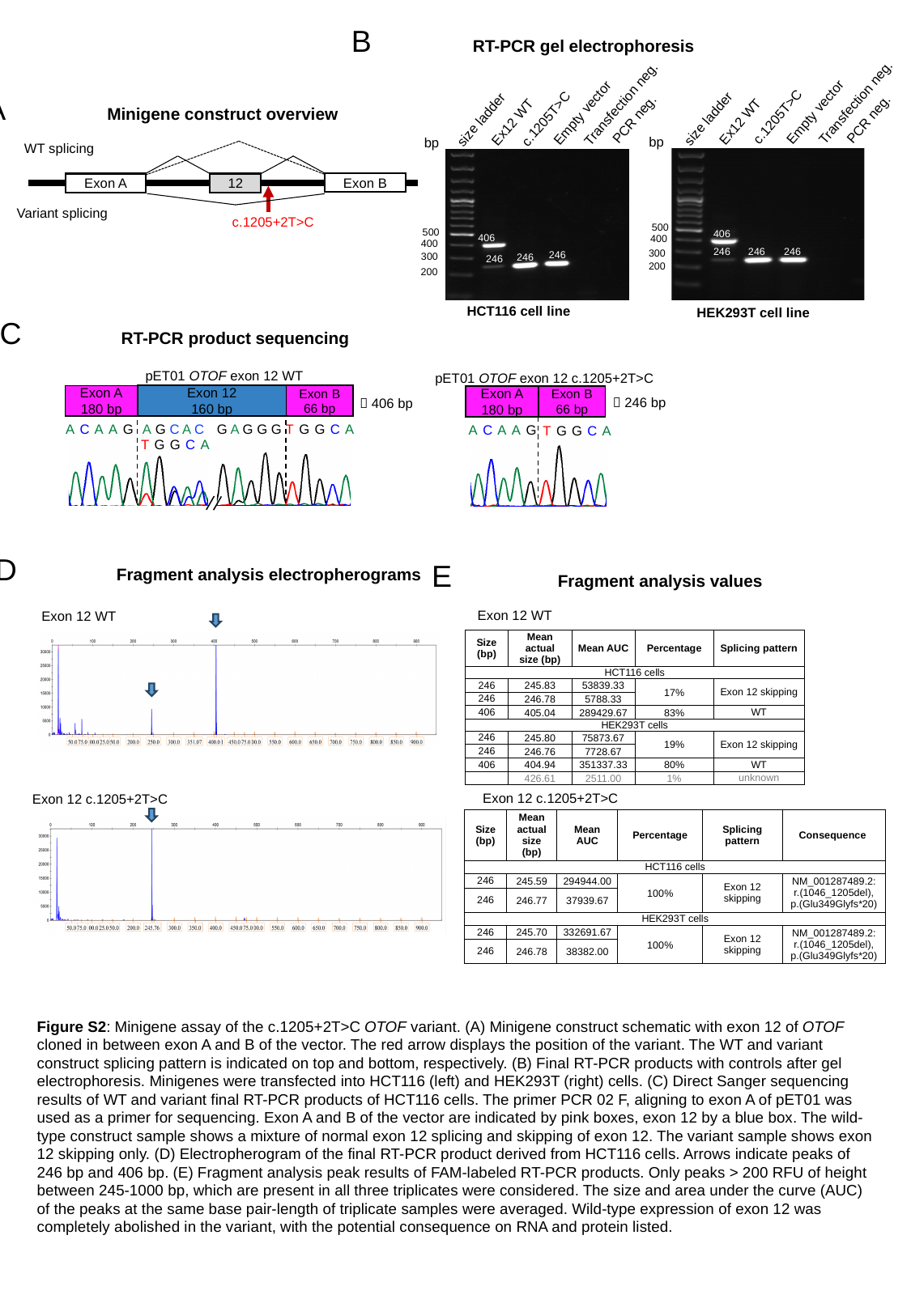

B	RT-PCR gel electrophoresis
c.1205T>C
 c.1205T>C
Transfection neg.
Transfection neg.
Ex12 WT
Ex12 WT
PCR neg.
PCR neg.
Empty vector
Empty vector
size ladder
size ladder
bp
bp
500
500
406
406
400
400
246
246
246
300
246
300
246
246
200
200
HCT116 cell line
HEK293T cell line
A	Minigene construct overview
WT splicing
Exon B
Exon A
12
Variant splicing
c.1205+2T>C
C	RT-PCR product sequencing
pET01 OTOF exon 12 WT
Exon 12
160 bp
Exon B
66 bp
Exon A
180 bp
ACAAG A G C A C
G A G G G TGGCA
TGGCA
pET01 OTOF exon 12 c.1205+2T>C
Exon A
180 bp
Exon B
66 bp
🡪 246 bp
ACAAG
TGGCA
🡪 406 bp
D	Fragment analysis electropherograms
E	 Fragment analysis values
Exon 12 WT
Exon 12 WT
Exon 12 c.1205+2T>C
| Size (bp) | Mean actual size (bp) | Mean AUC | Percentage | Splicing pattern |
| --- | --- | --- | --- | --- |
| HCT116 cells | | | | |
| 246 | 245.83 | 53839.33 | 17% | Exon 12 skipping |
| 246 | 246.78 | 5788.33 | | |
| 406 | 405.04 | 289429.67 | 83% | WT |
| HEK293T cells | | | | |
| 246 | 245.80 | 75873.67 | 19% | Exon 12 skipping |
| 246 | 246.76 | 7728.67 | | |
| 406 | 404.94 | 351337.33 | 80% | WT |
| | 426.61 | 2511.00 | 1% | unknown |
Exon 12 c.1205+2T>C
| Size (bp) | Mean actual size (bp) | Mean AUC | Percentage | Splicing pattern | Consequence |
| --- | --- | --- | --- | --- | --- |
| HCT116 cells | | | | | |
| 246 | 245.59 | 294944.00 | 100% | Exon 12 skipping | NM\_001287489.2: r.(1046\_1205del), p.(Glu349Glyfs\*20) |
| 246 | 246.77 | 37939.67 | | | |
| HEK293T cells | | | | | |
| 246 | 245.70 | 332691.67 | 100% | Exon 12 skipping | NM\_001287489.2: r.(1046\_1205del), p.(Glu349Glyfs\*20) |
| 246 | 246.78 | 38382.00 | | | |
Figure S2: Minigene assay of the c.1205+2T>C OTOF variant. (A) Minigene construct schematic with exon 12 of OTOF cloned in between exon A and B of the vector. The red arrow displays the position of the variant. The WT and variant construct splicing pattern is indicated on top and bottom, respectively. (B) Final RT-PCR products with controls after gel electrophoresis. Minigenes were transfected into HCT116 (left) and HEK293T (right) cells. (C) Direct Sanger sequencing results of WT and variant final RT-PCR products of HCT116 cells. The primer PCR 02 F, aligning to exon A of pET01 was used as a primer for sequencing. Exon A and B of the vector are indicated by pink boxes, exon 12 by a blue box. The wild-type construct sample shows a mixture of normal exon 12 splicing and skipping of exon 12. The variant sample shows exon 12 skipping only. (D) Electropherogram of the final RT-PCR product derived from HCT116 cells. Arrows indicate peaks of 246 bp and 406 bp. (E) Fragment analysis peak results of FAM-labeled RT-PCR products. Only peaks > 200 RFU of height between 245-1000 bp, which are present in all three triplicates were considered. The size and area under the curve (AUC) of the peaks at the same base pair-length of triplicate samples were averaged. Wild-type expression of exon 12 was completely abolished in the variant, with the potential consequence on RNA and protein listed.
