## Supplementary material for "Protocol for a minigene splice assay using the pET01 vector": Figure S3

### Slide 1
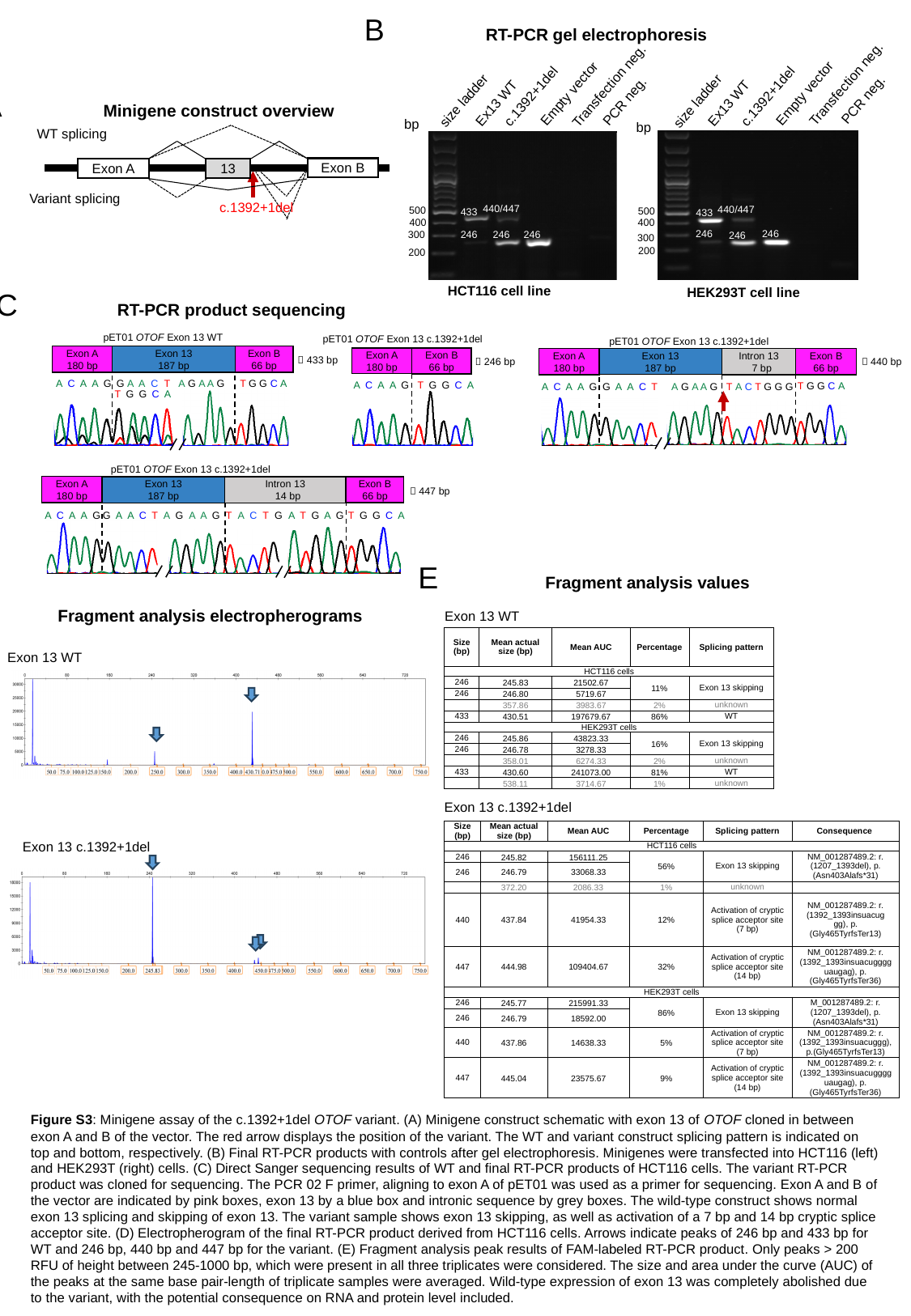

B	RT-PCR gel electrophoresis
c.1392+1del
c.1392+1del
Transfection neg.
Transfection neg.
Ex13 WT
Ex13 WT
PCR neg.
PCR neg.
Empty vector
Empty vector
size ladder
size ladder
bp
bp
500
500
406
406
400
400
246
246
300
246
246
246
300
246
200
200
HCT116 cell line
HEK293T cell line
440/447
440/447
433
433
246
246
246
246
246
246
A	Minigene construct overview
WT splicing
Exon B
Exon A
13
Variant splicing
c.1392+1del
C	RT-PCR product sequencing
pET01 OTOF Exon 13 WT
Exon 13
187 bp
Exon B
66 bp
Exon A
180 bp
🡪 433 bp
ACAAG
G A A C T
A G A A G
TGGCA
T G G C A
pET01 OTOF Exon 13 c.1392+1del
Exon A
180 bp
Exon B
66 bp
🡪 246 bp
TGGCA
ACAAG
pET01 OTOF Exon 13 c.1392+1del
Exon B
66 bp
Intron 13
 7 bp
Exon A
180 bp
Exon 13
187 bp
🡪 440 bp
🡪 bp
T G G C A
ACAAG
A G A A G T A C T G G G
GAACT
pET01 OTOF Exon 13 c.1392+1del
Exon A
180 bp
Exon B
66 bp
Exon 13
187 bp
Intron 13
 14 bp
🡪 447 bp
🡪 bp
TGGCA
ACAAG
ATGAG
A G A A G T A C T G
GAACT
E	 Fragment analysis values
D	 Fragment analysis electropherograms
Exon 13 WT
| Size (bp) | Mean actual size (bp) | Mean AUC | Percentage | Splicing pattern |
| --- | --- | --- | --- | --- |
| HCT116 cells | | | | |
| 246 | 245.83 | 21502.67 | 11% | Exon 13 skipping |
| 246 | 246.80 | 5719.67 | | |
| | 357.86 | 3983.67 | 2% | unknown |
| 433 | 430.51 | 197679.67 | 86% | WT |
| HEK293T cells | | | | |
| 246 | 245.86 | 43823.33 | 16% | Exon 13 skipping |
| 246 | 246.78 | 3278.33 | | |
| | 358.01 | 6274.33 | 2% | unknown |
| 433 | 430.60 | 241073.00 | 81% | WT |
| | 538.11 | 3714.67 | 1% | unknown |
Exon 13 WT
Exon 13 c.1392+1del
| Size (bp) | Mean actual size (bp) | Mean AUC | Percentage | Splicing pattern | Consequence |
| --- | --- | --- | --- | --- | --- |
| HCT116 cells | | | | | |
| 246 | 245.82 | 156111.25 | 56% | Exon 13 skipping | NM\_001287489.2: r.(1207\_1393del), p.(Asn403Alafs\*31) |
| 246 | 246.79 | 33068.33 | | | |
| | 372.20 | 2086.33 | 1% | unknown | |
| 440 | 437.84 | 41954.33 | 12% | Activation of cryptic splice acceptor site (7 bp) | NM\_001287489.2: r.(1392\_1393insuacuggg), p.(Gly465TyrfsTer13) |
| 447 | 444.98 | 109404.67 | 32% | Activation of cryptic splice acceptor site (14 bp) | NM\_001287489.2: r.(1392\_1393insuacugggguaugag), p.(Gly465TyrfsTer36) |
| HEK293T cells | | | | | |
| 246 | 245.77 | 215991.33 | 86% | Exon 13 skipping | M\_001287489.2: r.(1207\_1393del), p.(Asn403Alafs\*31) |
| 246 | 246.79 | 18592.00 | | | |
| 440 | 437.86 | 14638.33 | 5% | Activation of cryptic splice acceptor site (7 bp) | NM\_001287489.2: r.(1392\_1393insuacuggg), p.(Gly465TyrfsTer13) |
| 447 | 445.04 | 23575.67 | 9% | Activation of cryptic splice acceptor site (14 bp) | NM\_001287489.2: r.(1392\_1393insuacugggguaugag), p.(Gly465TyrfsTer36) |
Exon 13 c.1392+1del
Figure S3: Minigene assay of the c.1392+1del OTOF variant. (A) Minigene construct schematic with exon 13 of OTOF cloned in between exon A and B of the vector. The red arrow displays the position of the variant. The WT and variant construct splicing pattern is indicated on top and bottom, respectively. (B) Final RT-PCR products with controls after gel electrophoresis. Minigenes were transfected into HCT116 (left) and HEK293T (right) cells. (C) Direct Sanger sequencing results of WT and final RT-PCR products of HCT116 cells. The variant RT-PCR product was cloned for sequencing. The PCR 02 F primer, aligning to exon A of pET01 was used as a primer for sequencing. Exon A and B of the vector are indicated by pink boxes, exon 13 by a blue box and intronic sequence by grey boxes. The wild-type construct shows normal exon 13 splicing and skipping of exon 13. The variant sample shows exon 13 skipping, as well as activation of a 7 bp and 14 bp cryptic splice acceptor site. (D) Electropherogram of the final RT-PCR product derived from HCT116 cells. Arrows indicate peaks of 246 bp and 433 bp for WT and 246 bp, 440 bp and 447 bp for the variant. (E) Fragment analysis peak results of FAM-labeled RT-PCR product. Only peaks > 200 RFU of height between 245-1000 bp, which were present in all three triplicates were considered. The size and area under the curve (AUC) of the peaks at the same base pair-length of triplicate samples were averaged. Wild-type expression of exon 13 was completely abolished due to the variant, with the potential consequence on RNA and protein level included.
