## Supplementary material for "Protocol for a minigene splice assay using the pET01 vector": Figure S4

### Slide 1
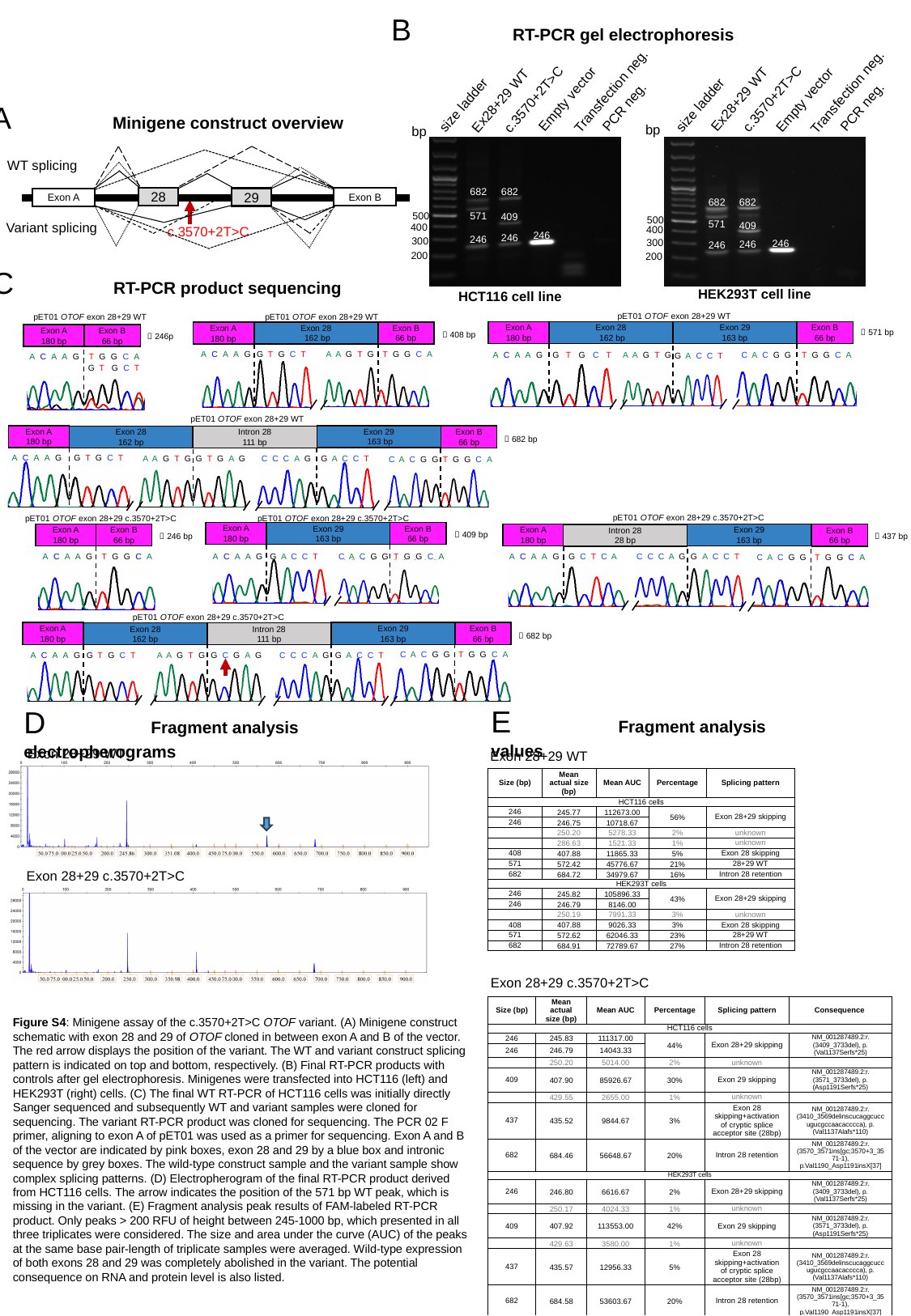

c.3570+2T>C
c.3570+2T>C
Transfection neg.
Transfection neg.
Ex28+29 WT
Ex28+29 WT
PCR neg.
PCR neg.
Empty vector
Empty vector
size ladder
size ladder
bp
bp
500
500
406
406
400
400
246
246
300
246
300
246
246
246
200
200
HEK293T cell line
HCT116 cell line
682
682
682
682
571
409
571
409
246
246
246
246
246
246
B	RT-PCR gel electrophoresis
A	Minigene construct overview
WT splicing
28
Exon B
Exon A
Variant splicing
c.3570+2T>C
29
C	RT-PCR product sequencing
pET01 OTOF exon 28+29 WT
🡪 571 bp
Exon B
66 bp
Exon A
180 bp
ACAAG
Exon 28
 162 bp
Exon 29
163 bp
A A G T G
G T G C T
C A C G G
T G G C A
G A C C T
pET01 OTOF exon 28+29 WT
Exon B
66 bp
Exon A
180 bp
🡪 408 bp
ACAAG
Exon 28
 162 bp
G T G C T
A A G T G
T G G C A
pET01 OTOF exon 28+29 WT
Exon B
66 bp
Exon A
180 bp
🡪 246p
ACAAG
TGGCA
G T G C T
pET01 OTOF exon 28+29 WT
Exon B
66 bp
Exon A
180 bp
🡪 682 bp
ACAAG
Exon 29
 163 bp
Exon 28
162 bp
Intron 28
111 bp
G T G C T
A A G T G
G A C C T
C A C G G
T G G C A
G T G A G
C C C A G
pET01 OTOF exon 28+29 c.3570+2T>C
Exon B
66 bp
Exon A
180 bp
ACAAG
Exon 29
163 bp
Intron 28
28 bp
C C C A G
G A C C T
G C T C A
TGGCA
C A C G G
🡪 437 bp
pET01 OTOF exon 28+29 c.3570+2T>C
Exon A
180 bp
Exon B
66 bp
🡪 246 bp
ACAAG
TGGCA
pET01 OTOF exon 28+29 c.3570+2T>C
Exon A
180 bp
Exon B
66 bp
🡪 409 bp
ACAAG
Exon 29
163 bp
G A C C T
C A C G G
TGGCA
pET01 OTOF exon 28+29 c.3570+2T>C
Exon B
66 bp
Exon A
180 bp
🡪 682 bp
ACAAG
Exon 29
163 bp
Exon 28
162 bp
Intron 28
111 bp
C A C G G
T G G C A
G T G C T
A A G T G
G C G A G
G A C C T
C C C A G
E	 Fragment analysis values
D	 Fragment analysis electropherograms
Exon 28+29 WT
Exon 28+29 WT
| Size (bp) | Mean actual size (bp) | Mean AUC | Percentage | Splicing pattern |
| --- | --- | --- | --- | --- |
| HCT116 cells | | | | |
| 246 | 245.77 | 112673.00 | 56% | Exon 28+29 skipping |
| 246 | 246.75 | 10718.67 | | |
| | 250.20 | 5278.33 | 2% | unknown |
| | 286.63 | 1521.33 | 1% | unknown |
| 408 | 407.88 | 11865.33 | 5% | Exon 28 skipping |
| 571 | 572.42 | 45776.67 | 21% | 28+29 WT |
| 682 | 684.72 | 34979.67 | 16% | Intron 28 retention |
| HEK293T cells | | | | |
| 246 | 245.82 | 105896.33 | 43% | Exon 28+29 skipping |
| 246 | 246.79 | 8146.00 | | |
| | 250.19 | 7991.33 | 3% | unknown |
| 408 | 407.88 | 9026.33 | 3% | Exon 28 skipping |
| 571 | 572.62 | 62046.33 | 23% | 28+29 WT |
| 682 | 684.91 | 72789.67 | 27% | Intron 28 retention |
Exon 28+29 c.3570+2T>C
Exon 28+29 c.3570+2T>C
| Size (bp) | Mean actual size (bp) | Mean AUC | Percentage | Splicing pattern | Consequence |
| --- | --- | --- | --- | --- | --- |
| HCT116 cells | | | | | |
| 246 | 245.83 | 111317.00 | 44% | Exon 28+29 skipping | NM\_001287489.2:r.(3409\_3733del), p.(Val1137Serfs\*25) |
| 246 | 246.79 | 14043.33 | | | |
| | 250.20 | 5014.00 | 2% | unknown | |
| 409 | 407.90 | 85926.67 | 30% | Exon 29 skipping | NM\_001287489.2:r.(3571\_3733del), p.(Asp1191Serfs\*25) |
| | 429.55 | 2655.00 | 1% | unknown | |
| 437 | 435.52 | 9844.67 | 3% | Exon 28 skipping+activation of cryptic splice acceptor site (28bp) | NM\_001287489.2:r.(3410\_3569delinscucaggcuccugucgccaacacccca), p.(Val1137Alafs\*110) |
| 682 | 684.46 | 56648.67 | 20% | Intron 28 retention | NM\_001287489.2:r.(3570\_3571ins[gc;3570+3\_3571-1), p.Val1190\_Asp1191insX[37] |
| HEK293T cells | | | | | |
| 246 | 246.80 | 6616.67 | 2% | Exon 28+29 skipping | NM\_001287489.2:r.(3409\_3733del), p.(Val1137Serfs\*25) |
| | 250.17 | 4024.33 | 1% | unknown | |
| 409 | 407.92 | 113553.00 | 42% | Exon 29 skipping | NM\_001287489.2:r.(3571\_3733del), p.(Asp1191Serfs\*25) |
| | 429.63 | 3580.00 | 1% | unknown | |
| 437 | 435.57 | 12956.33 | 5% | Exon 28 skipping+activation of cryptic splice acceptor site (28bp) | NM\_001287489.2:r.(3410\_3569delinscucaggcuccugucgccaacacccca), p.(Val1137Alafs\*110) |
| 682 | 684.58 | 53603.67 | 20% | Intron 28 retention | NM\_001287489.2:r.(3570\_3571ins[gc;3570+3\_3571-1), p.Val1190\_Asp1191insX[37] |
Figure S4: Minigene assay of the c.3570+2T>C OTOF variant. (A) Minigene construct schematic with exon 28 and 29 of OTOF cloned in between exon A and B of the vector. The red arrow displays the position of the variant. The WT and variant construct splicing pattern is indicated on top and bottom, respectively. (B) Final RT-PCR products with controls after gel electrophoresis. Minigenes were transfected into HCT116 (left) and HEK293T (right) cells. (C) The final WT RT-PCR of HCT116 cells was initially directly Sanger sequenced and subsequently WT and variant samples were cloned for sequencing. The variant RT-PCR product was cloned for sequencing. The PCR 02 F primer, aligning to exon A of pET01 was used as a primer for sequencing. Exon A and B of the vector are indicated by pink boxes, exon 28 and 29 by a blue box and intronic sequence by grey boxes. The wild-type construct sample and the variant sample show complex splicing patterns. (D) Electropherogram of the final RT-PCR product derived from HCT116 cells. The arrow indicates the position of the 571 bp WT peak, which is missing in the variant. (E) Fragment analysis peak results of FAM-labeled RT-PCR product. Only peaks > 200 RFU of height between 245-1000 bp, which presented in all three triplicates were considered. The size and area under the curve (AUC) of the peaks at the same base pair-length of triplicate samples were averaged. Wild-type expression of both exons 28 and 29 was completely abolished in the variant. The potential consequence on RNA and protein level is also listed.
