## Supplementary material for "Protocol for a minigene splice assay using the pET01 vector": Figure S5

### Slide 1
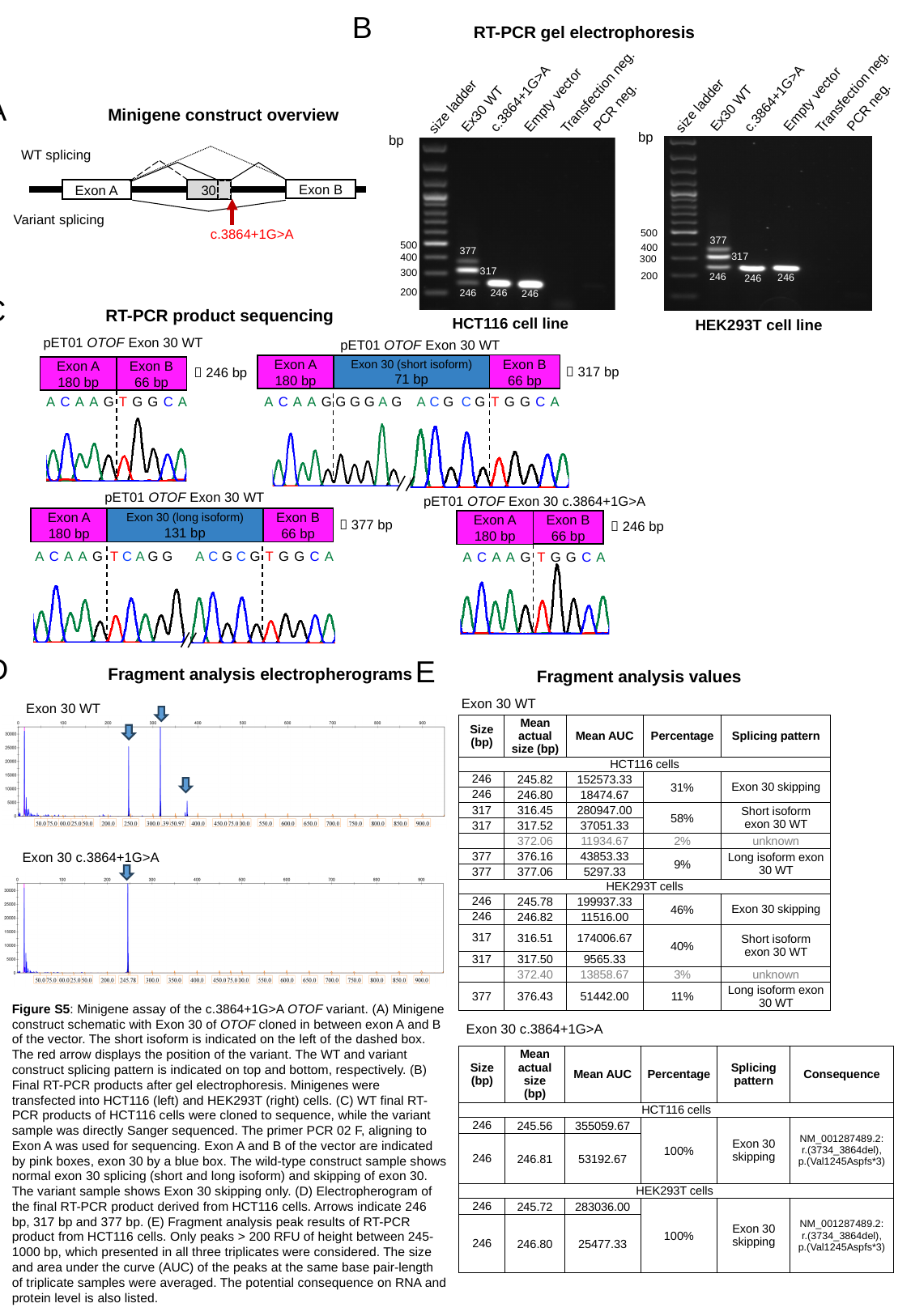

B	RT-PCR gel electrophoresis
c.3864+1G>A
Transfection neg.
 c.3864+1G>A
Transfection neg.
Ex30 WT
Ex30 WT
PCR neg.
PCR neg.
Empty vector
Empty vector
size ladder
size ladder
bp
bp
406
500
406
500
400
246
246
246
400
246
246
300
246
300
200
200
HCT116 cell line
HEK293T cell line
377
377
317
317
246
246
246
246
246
246
A	Minigene construct overview
WT splicing
Exon B
Exon A
30
Variant splicing
c.3864+1G>A
C	RT-PCR product sequencing
pET01 OTOF Exon 30 WT
Exon A
180 bp
Exon B
66 bp
🡪 246 bp
ACAAG
TGGCA
pET01 OTOF Exon 30 WT
Exon 30 (short isoform)
71 bp
Exon B
66 bp
Exon A
180 bp
🡪 317 bp
ACAAG
TGGCA
G G G A G
A C G C G
pET01 OTOF Exon 30 WT
Exon 30 (long isoform)
131 bp
Exon B
66 bp
Exon A
180 bp
🡪 377 bp
ACAAG
T C A G G
TGGCA
 A C G C G
pET01 OTOF Exon 30 c.3864+1G>A
Exon A
180 bp
Exon B
66 bp
🡪 246 bp
ACAAG
TGGCA
D	Fragment analysis electropherograms
Exon 30 WT
Exon 30 c.3864+1G>A
E	Fragment analysis values
Exon 30 WT
| Size (bp) | Mean actual size (bp) | Mean AUC | Percentage | Splicing pattern |
| --- | --- | --- | --- | --- |
| HCT116 cells | | | | |
| 246 | 245.82 | 152573.33 | 31% | Exon 30 skipping |
| 246 | 246.80 | 18474.67 | | |
| 317 | 316.45 | 280947.00 | 58% | Short isoform exon 30 WT |
| 317 | 317.52 | 37051.33 | | |
| | 372.06 | 11934.67 | 2% | unknown |
| 377 | 376.16 | 43853.33 | 9% | Long isoform exon 30 WT |
| 377 | 377.06 | 5297.33 | | |
| HEK293T cells | | | | |
| 246 | 245.78 | 199937.33 | 46% | Exon 30 skipping |
| 246 | 246.82 | 11516.00 | | |
| 317 | 316.51 | 174006.67 | 40% | Short isoform exon 30 WT |
| 317 | 317.50 | 9565.33 | | |
| | 372.40 | 13858.67 | 3% | unknown |
| 377 | 376.43 | 51442.00 | 11% | Long isoform exon 30 WT |
Figure S5: Minigene assay of the c.3864+1G>A OTOF variant. (A) Minigene construct schematic with Exon 30 of OTOF cloned in between exon A and B of the vector. The short isoform is indicated on the left of the dashed box. The red arrow displays the position of the variant. The WT and variant construct splicing pattern is indicated on top and bottom, respectively. (B) Final RT-PCR products after gel electrophoresis. Minigenes were transfected into HCT116 (left) and HEK293T (right) cells. (C) WT final RT-PCR products of HCT116 cells were cloned to sequence, while the variant sample was directly Sanger sequenced. The primer PCR 02 F, aligning to Exon A was used for sequencing. Exon A and B of the vector are indicated by pink boxes, exon 30 by a blue box. The wild-type construct sample shows normal exon 30 splicing (short and long isoform) and skipping of exon 30. The variant sample shows Exon 30 skipping only. (D) Electropherogram of the final RT-PCR product derived from HCT116 cells. Arrows indicate 246 bp, 317 bp and 377 bp. (E) Fragment analysis peak results of RT-PCR product from HCT116 cells. Only peaks > 200 RFU of height between 245-1000 bp, which presented in all three triplicates were considered. The size and area under the curve (AUC) of the peaks at the same base pair-length of triplicate samples were averaged. The potential consequence on RNA and protein level is also listed.
Exon 30 c.3864+1G>A
| Size (bp) | Mean actual size (bp) | Mean AUC | Percentage | Splicing pattern | Consequence |
| --- | --- | --- | --- | --- | --- |
| HCT116 cells | | | | | |
| 246 | 245.56 | 355059.67 | 100% | Exon 30 skipping | NM\_001287489.2: r.(3734\_3864del), p.(Val1245Aspfs\*3) |
| 246 | 246.81 | 53192.67 | | | |
| HEK293T cells | | | | | |
| 246 | 245.72 | 283036.00 | 100% | Exon 30 skipping | NM\_001287489.2: r.(3734\_3864del), p.(Val1245Aspfs\*3) |
| 246 | 246.80 | 25477.33 | | | |
