## Supplementary material for "Protocol for a minigene splice assay using the pET01 vector": Figure S6

**Supplementary Table S6:** Minigene assay primers

| Region of interest | Primer name | Primer sequence 5´- 3´ | Product size |
| --- | --- | --- | --- |
| Exon 12 | OTOF_Ex12 XhoI F | aattctcgagAGGGACCAAGACAGCATTTG | 454 bp |
|  | OTOF _Ex12 BamHI R | attggatccTTCTCAGCCATCCTCCATCA |  |
| Intron 12 c.1205+2T>C | OTOF_Ex12_ c.1205+2T-C_mut_F | CATTGAGGGGcGAGGCCCAGC | -- |
|  | OTOF_Ex12_ c.1205+2T-C_mut_R | TCATCTTCGTCGGTCTCATTGG |  |
| Exon 13 | hu_OTOF_Ex13_XhoI_F | aattctcgagTGTTCTTGGGAGGTGGGTAT | 553 bp |
|  | hu_OTOF_Ex13_BamHI_R | attggatccTGGCCAACATGATTTCCAG | -- |
| Intron 13 c.1392+1del | OTOF_Ex13_c.1392+1del_mut_F | TACTGGGGTATGAGGTAC |  |
|  | OTOF_Ex13_c.1392+1del_mut_R | CTTCTGGCCAGCAAAG |  |
| Exon 28 +29 | OTOF_Ex28+29 XhoI F | aattctcgagAATCTGGGGTGAACTACTGCC | 693 bp |
|  | OTOF _Ex28+29 BamHI R | attggatccTGACCTTCTTCTATGCCCACC |  |
| Intron 28 c.3570+2T>C | OTOF_c.3570+2T>C_Ex28_mut_F | TTTGAAGTGGcGAGTGCAGGC | -- |
|  | OTOF_c.3570+2T>C_Ex28_mut_R | CCACTTGACGAGGGTG |  |
| Exon 30 | hu_OTOF_Ex30_XhoI_F | aattctcgagCGCTGGTTGATGGAGAAGA | 676 bp |
|  | hu_OTOF_Ex30_BamHI_R | attggatccATACGCATGCACGCACAA |  |
| Intron 30 c.3864+1G>A | OTOF_c.3864+1G>A_Ex30_mut_F_2^nd^ | GCTGGACGCGaTAAGGCGGGT | -- |
|  | OTOF_c.3864+1G>A_Ex30_mut_R_2^nd^ | TTCACCATGGTCTCCAGTTTCTTGATGG |  |
